# Culture-supported ecophysiology of the SAR116 clade demonstrates metabolic and spatial niche partitioning

**DOI:** 10.1101/2025.01.06.631592

**Authors:** Jordan T. Coelho, Lauren Teubner, Michael W. Henson, V. Celeste Lanclos, Conner Y. Kojima, J. Cameron Thrash

## Abstract

Marine SAR116 bacterioplankton are ubiquitous in surface waters across global oceans and form their own order, *Puniceispirillales,* within the *Alphaproteobacteria.* To date no comparative physiology among diverse SAR116 isolates has been performed to capture the functional diversity within the clade, and further, diversity through the lens of metabolic potential and environmental preferences via clade-wide pangenomics remains poorly constrained. Using high-throughput dilution-to-extinction cultivation, we isolated and genome sequenced five new and diverse SAR116 isolates from the northern Gulf of Mexico. Here we present a comparative physiological analysis of these SAR116 isolates, along with a pangenomic investigation of the SAR116 clade using a combination of metagenome-assembled genomes (MAGs, n=258), single-amplified genomes (SAGs, n=84), previously existing (n=2), and new isolate genomes (n=5), totaling 349 SAR116 genomes. Phylogenomic investigation supported the division of SAR116 into three distinct subclades, each with additional structure totalling 15 monophyletic groups. Our SAR116 isolates belonged to three groups within subclade I representing distinct genera with different morphologies and varied phenotypic responses to salinity and temperature. Overall, SAR116 genomes encoded differences in vitamin and amino acid synthesis, trace metal transport, and osmolyte synthesis and transport. They also had genetic potential for diverse sulfur oxidation metabolisms, placing SAR116 at the confluence of the organic and inorganic sulfur pools. SAR116 subclades showed distinct patterns in habitat preferences across open ocean, coastal, and estuarine environments, and three of our isolates represented the most abundant coastal and estuarine subclade. This investigation provides the most comprehensive exploration of SAR116 to date anchored by new culture genomes and physiology.

## Introduction

Marine SAR116 bacterioplankton, first discovered in the Sargasso Sea [1], are ubiquitous across the epipelagic global oceans [2–4]. While peak relative abundances may reach up to 20% of the prokaryotic community during a summer bloom [5], they are more commonly found to represent 5-12% of the prokaryotic community [2, 5–9]. SAR116 are frequently associated with coastal phytoplankton blooms and tend to increase in abundance mid- to late-bloom [10–12]. In addition, SAR116 have been detected among marine sponge and coral samples [13–15], though the phylogenetic relationship of coral and sponge representatives of SAR116 has yet to be resolved. Early investigations into the evolutionary history of SAR116 delineated them as a monophyletic group within the *Alphaproteobacteria* [2, 5, 9, 13, 16], and although they were previously described as members of the *Rhodospirillaceae* [17, 18], more recently they have been classified in their own order, the *Puniceispirillales* [19, 20].

Advancements in the cultivation of marine microorganisms via high-throughput, dilution-to-extinction culturing using natural seawater media [21–23] lead to the isolation of the first three SAR116 cultured representatives [17, 18, 24], defined their aerobic chemoheterotrophic metabolism, and facilitated further genomic and physiological characterizations of the clade [17, 18]. Although extensive physiological characterizations of HTCC8037 [24] and HIMB100 [18] have yet to be published, strain IMCC1322 has received more experimental attention. Its genome encodes a green-tuned proteorhodopsin (522 nm) and light exposure enabled growth in a 20% (v/v) CO headspace compared with dark incubations where cells experienced CO toxicity [13]. More recently, IMCC1322 was shown to constitutively express proteorhodopsin regardless of the presence or absence of light [25].

Numerous investigations of *in situ* microbial assemblages have generated additional information on SAR116 physiology and spatial distributions. SAR116 have been described as active members of coastal and open-ocean systems [12] with particularly high transcriptional activity in coastal [26, 27] and estuarine waters [28]. In fact, at the Sapelo Island Microbial Observatory estuarine site, the IMCC1322 genome recruited the most transcripts of the sixteen diverse references used, implicating similar SAR116 taxa in polyphosphate and nitrate processing, and demonstrating the highest expression in genes for amino acids and five-carbon carbohydrates than other bacterioplankton [26]. In the Columbia River estuary, SAR116 were most abundant in samples with brackish salinities of 15.4 and 25.4, with high expression of metalloendopeptidases, and zinc, iron, and nickel transporters [28]. Among bacterioplankton with dimethylsulfoniopropionate (DMSP) metabolism via DMSP lyase (*dddP*) [29], two *dddP* gene operational taxonomic units (OTUs) belonging to SAR116 (OTU43 and OTU28) were the most abundant taxa, comprising 82 ± 14% of *dddP* genes in the upper euphotic zone across the NW Pacific transect [29]. This predicted physiology was experimentally validated in IMCC1322, which produced dimethyl sulfide (DMS) when supplemented with DMSP, a hallmark of DMSP lyase activity [29]. Of note, OTU43 was abundant and ubiquitous across the coastal and oligotrophic open-ocean sampling sites, while OTU28 was only detected in the oligotrophic open-ocean sites, suggesting biogeographic differentiation within the SAR116 clade. Overall, these *in situ* findings illuminated additional SAR116 physiologies and suggested specialized spatial distributions, indicating a much wider breadth of functional and ecological diversity than what was initially known from the first two isolates.

A previous clade-wide comparative genomic analysis classified the *Puniceispirillales* into two discrete subclades with different evolutionary histories [30]. Genomic characteristics such as GC content and intergenic spacer length suggested that one of these subclades had undergone streamlining [30] that evolutionarily selects for reductions in genome size and cell complexity [31]. This streamlined subclade was also more globally distributed across the TARA Oceans dataset [32], and its success was attributed to adaptation for a broader range of conditions relative to the non-streamlined SAR116 subclade [30]. This runs counter to canonical genome streamlining theory that predicts evolution towards a specialist rather than a generalist phenotype [31, 33, 34]. Nevertheless, these data demonstrated large-scale evolutionary and habitat heterogeneity within the SAR116 clade, and generated many new questions about the life history and microdiversity within this group.

Our recent isolation of five new SAR116 representatives [7], coupled to the reconstruction of hundreds of new SAR116 metagenome-assembled genomes (MAGs) [35], offered an opportunity to explore physiological diversity and update previous observations of the *Puniceispirillales*, especially related to the known boundaries of SAR116 functional diversity. To contextualize our isolates within the *Puniceispirillales*, and to more comprehensively investigate the functional diversity and niche partitioning across the clade, we employed comparative physiology, microscopy, and genomics with new and previously sequenced SAR116 isolate genomes (n=5 and n=2, respectively), MAGs (n=258), and single-amplified genomes (SAGs, n=84) for a total of 349 *Puniceispirillales* genomes. Here we have found the SAR116 clade to be composed of three distinct subclades with further subclade structure leading to 15 subclades in total. We predict that differentiation among subclades was driven by habitat preferences and metabolic niche partitioning, including osmolyte synthesis and transport, vitamin and amino acid synthesis, and sulfur oxidation metabolisms. Physiological characterization of our SAR116 isolates demonstrated their salinity preferences, along with phenotypic variation in temperature ranges and cell morphologies. These results expand our understanding of *Puniceispirillales* functional diversity and enable us to better constrain the ecological niches of subgroups within the SAR116 clade.

## Methods

### Isolation, genome sequencing, and assembly

We isolated new SAR116 strains using high-throughput dilution-to-extinction cultivation and identified them via 16S rRNA gene comparisons as described [7]. Genomic DNA sequencing and library preparation were completed at the University of Southern California (USC) Molecular Genomics Core using Illumina NextSeq 550. Additionally, we performed Oxford Nanopore long-read sequencing in house on four of the five genomes. We generated hybrid assemblies with Unicycler v0.4.8 [36], and used SPAdes v3.13.1 [37] for our Illumina-only assembly. Assembled genomes were assessed for quality and completeness using CheckM [38]. Detailed methods on sequencing, quality checking, and assembly, are in <u>Supplemental Text</u> [37, 39, 40].

### Additional Taxon Selection

We combined our five SAR116 isolate genomes with all publicly available SAR116 genomes from the Genome Taxonomy Database (GTDB, release89) [19, 20], Global Ocean Reference Genomes Tropics (GORG-Tropics) [41], and the OceanDNA MAG catalog [35]. Quality assessment, completion, and genome dereplication were done as previously described [42, 43]. `We required all genomes to have < 5% contamination, MAGs to have ≥ 70% completion, and SAGs to have ≥ 50% completion to remain in the analysis, resulting in a total of 349 SAR116 genomes (`Table S1). Outgroup taxa (n=31) for phylogenomic analysis were selected from cultured representatives across various orders of *Alphaproteobacteria* as described in Muñoz-Gómez et al. 2019 [44] and assessed for quality, completeness, and dereplicated as described above.

### Phylogenomics

We completed phylogenomic analysis in Anvi’o [45] v7.1 as previously described [42, 43]. Amino acid sequences were retrieved using ‘anvi-get-sequences-for-hmm-hits’ using the ‘--Rinke_et_al’ HMM profile [46]. Genes were trimmed with TrimAl [47] and aligned with MUSCLE [48], both with default settings, then concatenated using geneStitcher [49]. We used IQ-TREE v2.1.2 [50] for maximum-likelihood inference using traditional bootstrapping with 1000 replicates, and the automated amino acid substitution best-fit model estimator ‘-m MFP’ which selected LG+R10 as the best model. We visualized the resulting tree using *Ggtree* v3.2.1 and *Treeio* v1.18.1 R packages [51–53], rooted at the midpoint, and nodes were ordered in increasing order.

### Pangenomics

We conducted pangenomic analysis in Anvi’o [45] v7.1 similarly as reported [43], generating the pangenome annotation summary using *anvi-summarize,* and annotation lists for sulfur oxidation analyses can be found in Tables S2. Additionally, we used KEGG Decoder and Expander [54] to identify diversity in physiological and metabolic gene pathways, modified to include the osmolyte table in Henson et al. [55], along with additional genes associated with vitamin, organic carbon, amino acid, and metal transport (Table S3). With the output data we predicted subclade physiologies by calculating the median pathway completion values for each subclade. We visualized the output data using custom R-Scripts [56, 57]. Metagenomic recruitment was done as previously described [42, 43]. Reads-Per-Kilobase-Mapped (RPKM) was used as a proxy for subclade abundance and calculated using RRAP [58]. Additional details on pangenomics in <u>Supplemental Text</u> [59–65].

### Single-gene phylogenetics

We generated a 16S rRNA gene phylogeny comparison with previous 16S rRNA subclade designations [13], and to investigate the coral and sponge association among SAR116 members [66]. Sulfur oxidation gene phylogenies were also generated to evaluate the evolutionary histories of these proteins within SAR116. Additional details in <u>Supplemental Text</u> [67–75].

### Temperature and salinity tolerances, and environmental linear regressions

To test the temperature and salinity tolerances of cultured representatives, three LSUCC SAR116 isolates – LSUCC0719, LSUCC0744, and LSUCC0684 - were inoculated from 1 mL cryostocks into 5 mL of our complex and defined artificial seawater medium, MWH2 [76] using 10 mL sterile borosilicate glass test-tubes (#1512, Globe Scientific, New Jersey, USA). A range of seven temperatures and five salinities were tested. Growth for all experiments was measured with a BD Accuri C6 Plus flow cytometer (BD, New Jersey, USA) using 1x SYBR Green as described [7, 42] and plotted using sparse-growth-curve [77]. Additional details on experimental design in <u>Supplemental Text</u>.

### Scanning electron microscopy

We inoculated three LSUCC SAR116 isolates, LSUCC0719, LSUCC0744, and LSUCC0684, from cryostocks (1 mL) into 100 mL of MWH2 medium [76] in sterile 125 mL polycarbonate flasks (FPC0125S, TriForest, Irvine, CA), and fixed them at a final concentration of 2.5% (v/v) glutaraldehyde (G5882, Sigma-Aldrich) after growth to mid-exponential phase. Scanning electron microscopy (SEM) was performed at the USC NanoImaging Center, and SEM images were analyzed in ImageJ [78] to quantify cell size measurements. Additional details on SEM image prep and SEM image analysis in <u>Supplemental Text</u>.

## Results

### New genome characteristics

We sequenced and assembled five new SAR116 genomes spanning a wide diversity within subclade I (Figure 1a). Four out of five isolate assemblies produced closed, circular genomes, and all had 0% estimated contamination (Table 1). CheckM estimated the genome of LSUCC0684 to be 97.98% complete, however the genome assembled into a single circular contig, and thus the LSUCC0684 genome invites re-evaluation of the appropriate single-copy marker genes designated for SAR116 in CheckM. The LSUCC isolate genomes ranged in size from 2.59 – 2.80 Mbp with a G+C content range of 51% – 58%, and a predicted protein coding gene range of 2,448 – 2,658. The full set of SAR116 genomes all encoded a single rRNA operon and had estimated genome sizes ranging from 1.76 – 3.95 Mbp (mean 2.25 Mbp) (Figure 1b), G+C contents spanning 29 – 63% (Figure 1c), and coding densities from 81 – 95.7% (Figure 1d), all of which represented extremely large intraclade ranges.

**Figure 1:**
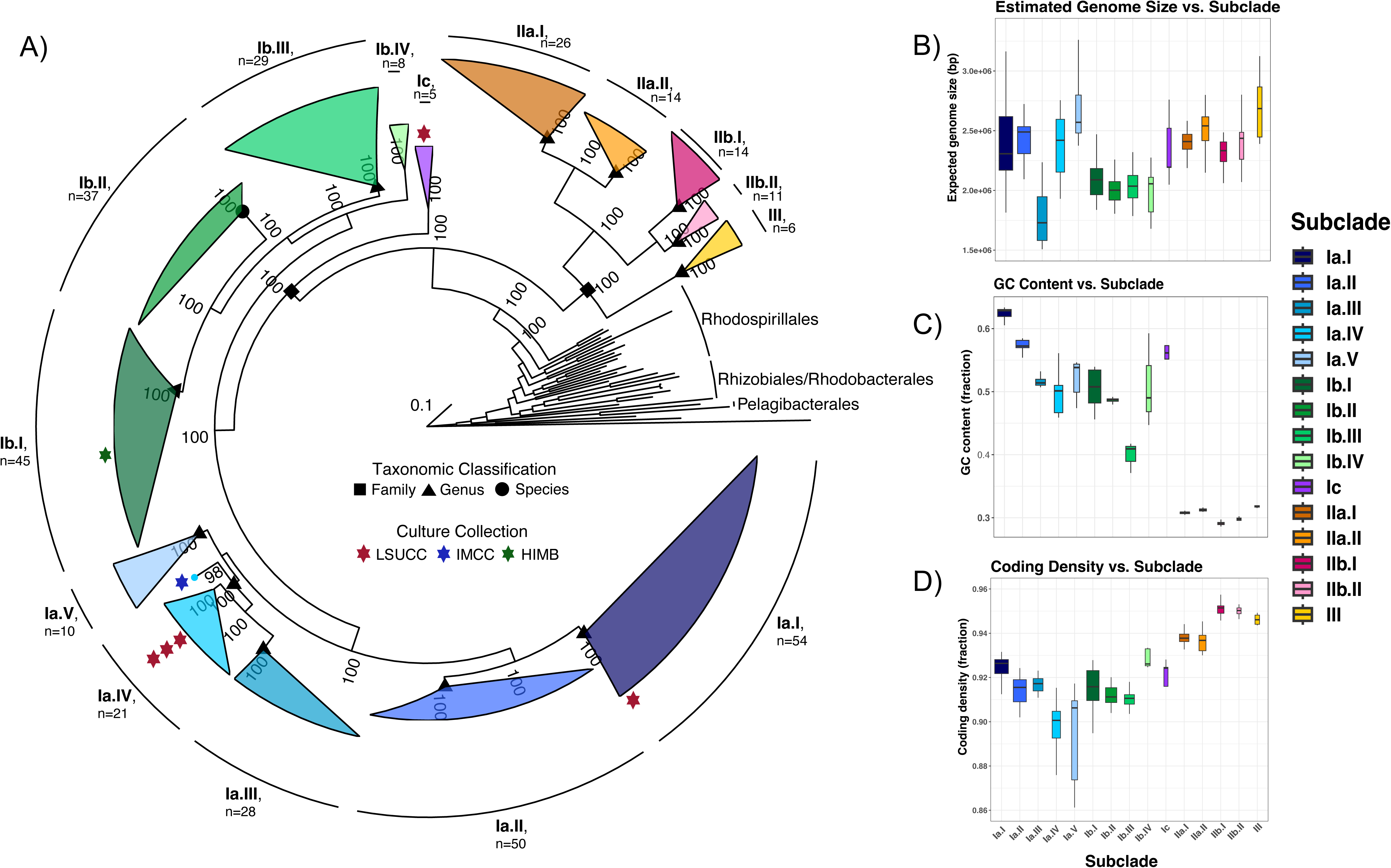
Subclade structure and genome characteristics across the SAR116 clade. (A) Phylogenomic tree of the SAR116 clade with other members of the *Alphaproteobacteria* as an outgroup. The scale bar represents 0.1 changes per position. Bootstrap support values (n=1000) are indicated at nodes. Red stars next to subclades indicate isolate representation and the color designates the culture collection the isolate is housed in according to the key inside the tree. Shapes overlaying internal nodes of SAR116 subclades represent the taxonomic classification of the descendants in the subclade according to the key inside the tree. Panels B-D indicate the estimated genome size (B), GC content (C), and coding density (D) (estimated via CheckM) of SAR116 genomes by subclade.

**Table 1:** Assembly statistics for new LSUCC SAR116 isolate representative genomes generated in this study. Genome size estimates were calculated by dividing the assembly size by the estimated percent completion (fraction) output from CheckM.

|  | LSUCC0719 | LSUCC0226 | LSUCC0396 | LSUCC0744 | LSUCC0684 |
| --- | --- | --- | --- | --- | --- |
| **Completion\*** | 99.13% | 100% | 100% | 100% | 100% |
| **Contamination** | 0.00% | 0.00% | 0.00% | 0.00% | 0.00% |
| **Actual/Est. Size** | 2.80 Mbp | 2.68 Mbp | 2.59 Mbp | 2.65 Mbp | 2.70 Mbp |
| **N50** | 535,962 | 2,678,285 | 2,592,803 | 2,653,851 | 2,704,759 |
| **Contigs Assembled** | 43 | 1 | 1 | 1 | 1 |

|  | LSUCC0719 | LSUCC0226 | LSUCC0396 | LSUCC0744 | LSUCC0684 | Other SAR116 |
| --- | --- | --- | --- | --- | --- | --- |
| **Completion\*** | 99.13% | 100% | 100% | 100% | 100% | 50.74 - 99.94%<br>*Avg: 79.98%* |
| **Contamination** | 0.00% | 0.00% | 0.00% | 0.00% | 0.00% | 0.00 - 4.99%<br>*Avg: 1.92%* |
| **Actual/Est. Size** | 2,796,311 bp | 2,678,285 bp | 2,592,803 bp | 2,635,851 bp | 2,704,644 bp | 1.51 - 3.95 Mbp<br>*Avg: 2.25 Mbp* |
| **Contigs Assembled** | 43<br>*N50: 535,962* | 1<br>*N50: 2,678,285* | 1<br>*N50: 2,592,803* | 1<br>*N50: 2,635,851* | 1<br>*N50: 2,704,644* | 1 - 1048<br>*Avg # contigs: 363* |
| **GC Content (fraction)** | 0.58 | 0.51 | 0.51 | 0.51 | 0.55 | 0.29 - 0.63<br>*Avg: 0.49* |
| **Predicted Genes** | 2658 | 2536 | 2428 | 2485 | 2551 | 1055 - 2875<br>*Avg: 1999* |
| **rRNA Operon Copy Number** | 1 | 1 | 1 | 1 | 1 | 1 |

### Phylogenetic diversity

Our updated phylogeny, using 380 total genomes (SAR116 n=349, Outgroup n=31) and average amino acid identity (AAI), supported dividing the SAR116 clade into three major subclades (I, II, and III) with additional structure defining 15 subclades in total (Fig. 1a, Figure S1, Table S4). This updated phylogeny recovered the same major subclade structure established via 16S rRNA genes [13] (Figure 2, S2), and a similar structure to a previous phylogenomic inference [30], however we have defined a third major subclade and have also expanded on the known diversity of subclade I, previously denoted as the high GC group (HGC) [30]. Here, our updated subclades II and III comprise the previously defined low GC (LGC) subclade [30]. 16S rRNA gene identity between the two most divergent genomes in the SAR116 clade (Figure S1) was 86.98% via BLAST [61], and pairwise AAI was 52.47%, both supported an Order level divergence [62], matching the GTBD classification. Roda-Garcia et al. proposed *Puniceispirillales* to comprise four families [30], however we conservatively confirmed the presence of two families using AAI taxonomic classification [63]. One family corresponds to subclade I and the other to subclades II and III (Figure 1a). Furthermore, we assigned putative genus demarcations to 12 of the 15 subclades (Figure 1a). The newly assembled genomes LSUCC0226, LSUCC0396, and LSUCC0744 branched within subclade Ia.IV, LSUCC0719 in subclade Ia.I, and LSUCC0684 in subclade Ic (Figure 1a). Thus, our five new isolates belonged to three separate genera that spanned the diversity of subclade I.

**Figure 2:**
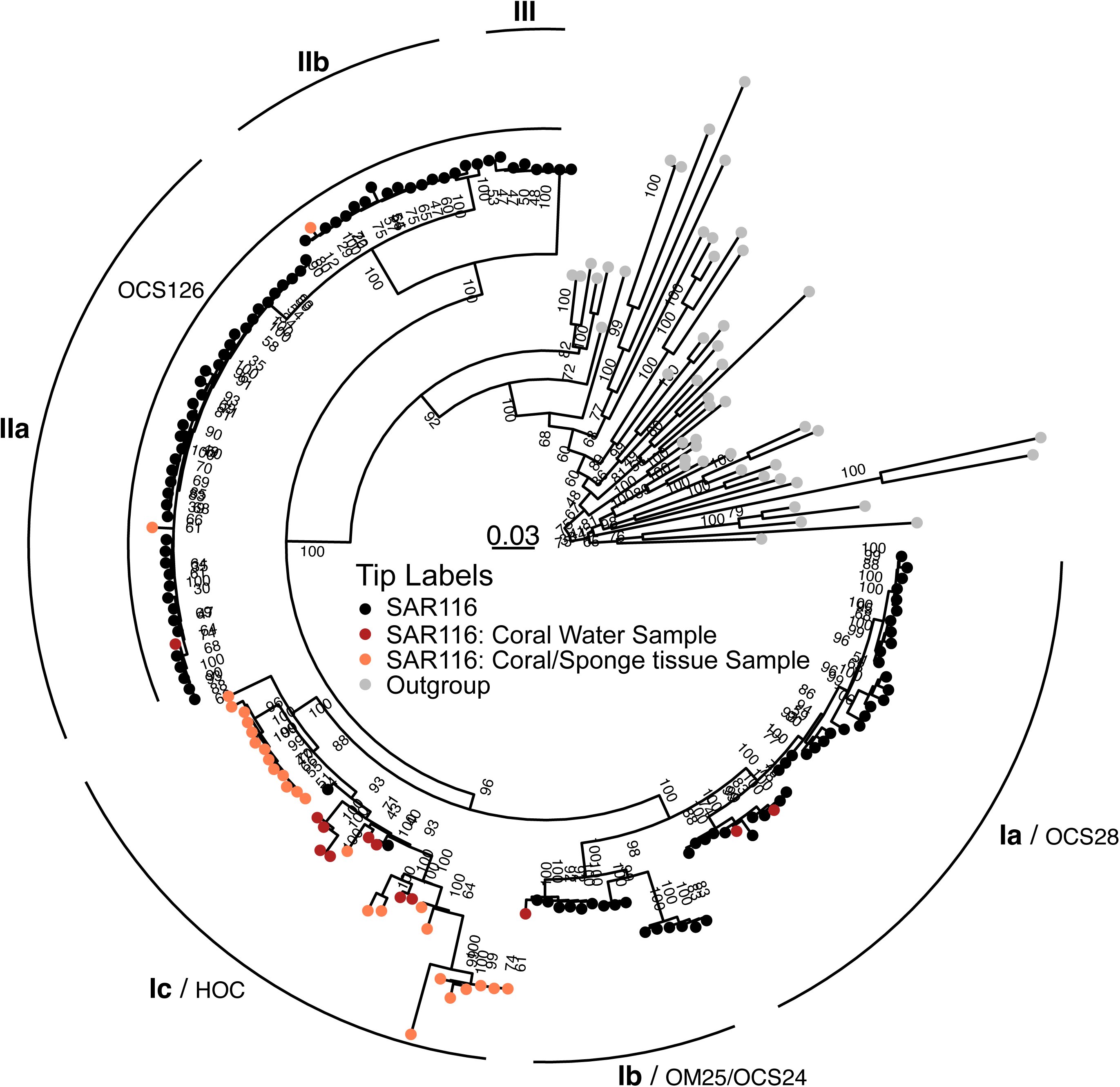
16S rRNA gene phylogenetic tree of SAR116 members and outgroup members of the *Alphaproteobacteria*. The scale bar represents 0.3 changes per position. Bootstrap values (n=1000) are represented at nodes. Black tip labels indicate SAR116 members, red tip labels indicate SAR116 members from water sampled at the coral:water interface, orange tip labels indicate SAR116 members from ground coral tissue samples, and gray tip labels indicate outgroup members. Subclades are labeled with the updated subclade designations from this investigation (left) and also the historical 16S subclade designations (right).

### Host-association

Although SAR116 is a cosmopolitan marine surface group [79–81], it has also been previously detected in marine sponge and coral tissues [14, 15, 66, 82], with relative abundances up to 10% of the prokaryotic endosymbiont community [66]. We investigated subclade affiliations of the putative endosymbiont SAR116 members in the context of our revised taxonomy to understand evolutionary origins of these taxa. The overwhelming majority of coral/sponge-associated SAR116 16S rRNA gene sequences grouped with subclade Ic, with only two host-associated sequences branching in subclade II (Figure 2). This suggests a specific endosymbiotic host-associated divergence within the *Puniceispirillales*. However, the presence of non-host associated sequences in the Ic subclade, one from the LSUCC0684 isolate genome and the other from the AG-917-F20 SAG collected at the Bermuda Atlantic Time Series (BATS) [41], supports the assertion that host-associated *Puniceispirillales* have a planktonic life stage [82]. Regardless, the current evidence suggests that host-association does not occur commonly within the SAR116 clade, and is primarily restricted to subclade Ic.

### New Metabolic Observations

We compared gene content in SAR116 to delineate differences in metabolic potential across subclades. Broadly, we recapitulated previous observations that *Puniceispirillales* genomes have predicted capacity for gluconeogenesis, the TCA cycle, and incomplete glycolysis (via Embden-Meyerhoff-Parnas) pathways [17, 18, 30] (Figure 3). And, similar to previous results [13, 17, 18] carbon monoxide oxidation genes (*coxSML)* were present throughout the SAR116 clade (Table S5). Proterhodopsin, the light-mediated proton-pump, has been previously described in SAR116 [13]. Here we found it to be prolific across the clade, however in our updated phylogeny, subclade Ia.III does not encode for the protein (Figure 3). There was no genetic evidence of carbon fixation or carbon dioxide assimilation, nor was there evidence of anaerobic respiration, thus supporting previous findings of an obligate aerobic and heterotrophic physiology [13, 18, 30]. Below are our predictions of SAR116 metabolic potential that have changed based on our updated analysis with new closed isolate genomes, and/or have not been previously discussed in the most recent SAR116 pangenomics analysis [30].

**Figure 3:**
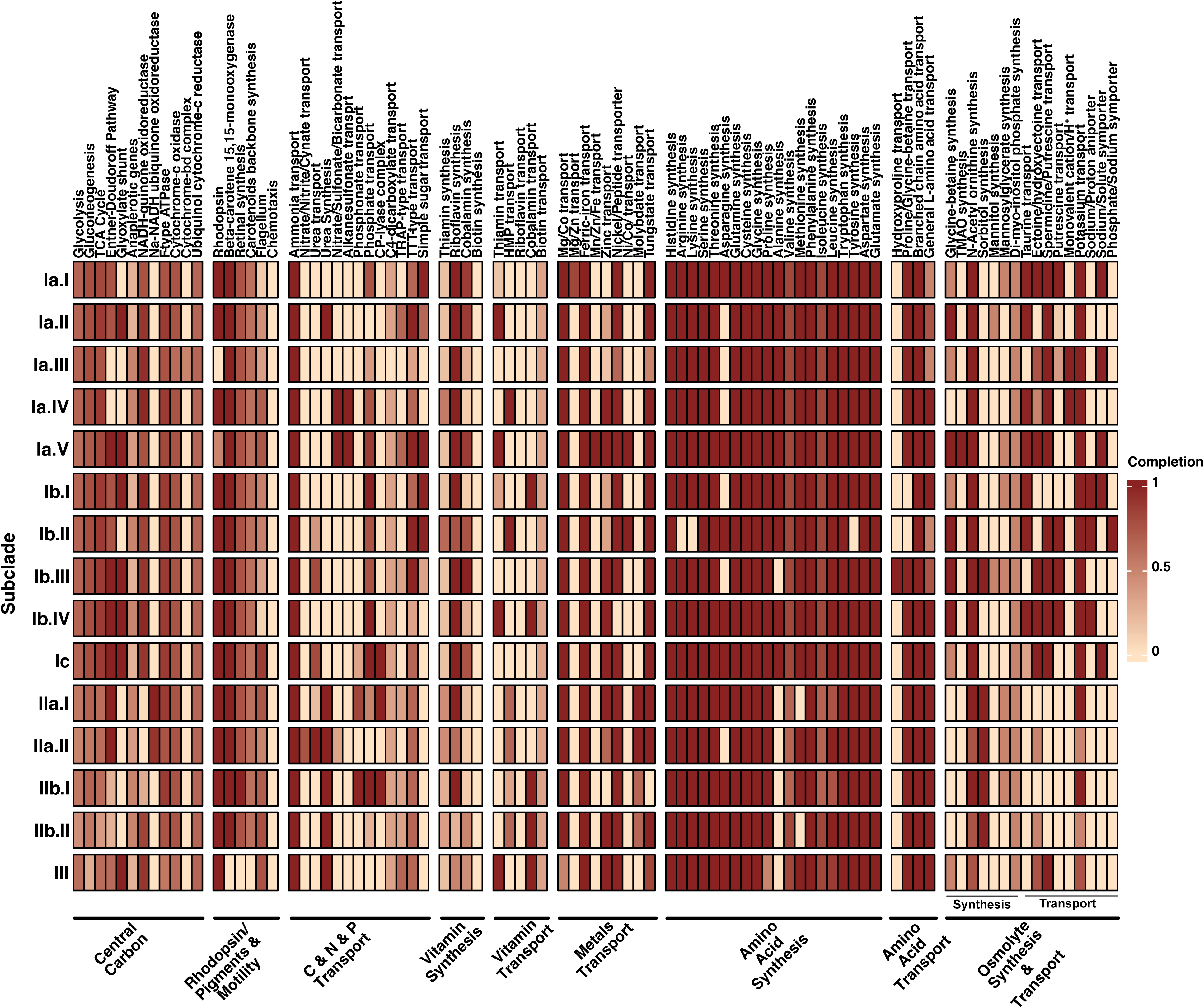
Output from KEGG-Decoder modified to include osmolyte synthesis and transport genes along with additional metals and vitamin transporters (Table S3). Each row is the median pathway representation by subclade, and every column is a metabolic pathway. The darker the color the more complete the pathway, with the pale peach color indicating no detection and dark red indicating a complete pathway.

Trace metal transport potential was diverse across the SAR116 clade, with few systems conserved in all subclades (Figure 3). We predicted that SAR116 predominantly rely on ferric iron transport through the *afuABC* complex to meet their iron needs. One subclade, Ia.V, carries a manganese/zinc/iron transporter (*sitABCD)*, that may also transport ferrous iron. Other metals that appeared important based on widely conserved transporters were magnesium and/or cobalt (all but subclade III encoded for a magnesium/cobalt transporter, *corC*), and tungsten (via *tupABC* is present in all subclades except IIb.I). Nickel may also be important for SAR116. The nickel/cobalt transporter, *rcnA*, was not widely distributed (only in subclades Ia.V and Ib.II). However, all but subclade Ib.IV had the nickel/peptide transporter, *ABC.PE.SPP1* (Figure 3), so subclades may obtain nickel via different systems. More sparsely distributed were transporters for zinc and manganese (subclades Ia.IV - V, Ib.III - IV, Ic, Ia.I, Ib.II, and III had *znuABC*; only subclade Ia.I had the manganese/zinc transporter *ABC.ZM.SAP*), as well as molybdate (*modABC* was only in subclade II) (Figure 3). While it remains possible that the absence of some of these transporters may result from the lack of complete genomes, the cases where a transporter was missing in all genomes from a subclade likely indicates a true gene absence. The differential distributions of these metal transporters across SAR116 offer potential insight into how the subclades are interacting with metals, and how trace metal scarcity can limit metalloenzyme activity and overall physiology. Namely, subclade Ia.V enocodes the most metal transporters of all SAR116 subclades, possibly indicating a diverse trace metals requirement among their proteome relative to the other SAR116 subclades. Additionally, while TupABC has a higher affinity for tungstate, it also transports molybdate [83]; thus the supplementary molybdate transport, *modABC*, unique to subclade II suggests an elevated requirement for molybdate in that group.

Vitamin synthesis and transport was also quite differentiated across the SAR116 subclades (Figure 3). Most subclades contained genomes with evidence of riboflavin synthesis, and several subclades had genomes with genes for cobalamin synthesis. Some subclades that lacked complete cobalamin synthesis appeared to compensate with cobalamin transport (Ib.I, Ib.IV, II.bI, II.bII, III). Many subclades had genes either for thiamin or the thiamin precursor, 4-amino-5-hydroxymethyl-2-methylpyrimidine (HMP), but not both. Some had neither (Figure 3). We did not find evidence of biotin synthesis genes in most genomes, but there were potential biotin transporters in most subclades.

SAR116 subclades were predicted to make most amino acids, with some notable exceptions (Figure 3). We did not find evidence for alanine synthesis in subclade Ib.III genomes, nor in any genomes of subclades II and III. Similarly, we did not find genes for methionine synthesis in subclade IIb.II; and while a little more than half of the genomes in subclade IIa.I did not carry genes for methionine synthesis, the remaining genomes were prototrophic for methionine (Figure S3). Given an average genome completion of 70% for subclade IIa.I (Figure S4), this inconsistency may be due to incomplete genomes. We also predicted asparagine synthesis to be missing completely from Ia.II and Ia.III, as asparagine synthesis genes were not detected across any genomes within these subclades (Figure S3). However in subclades Ia.IV, Ib.I, Ib.III, and IIa.II, there were asparagine synthesis pathways speckled across genomes in these subclades (Figure S3). While this may be due to incomplete genomes, even the closed isolate genomes in subclade Ia.IV reflected the inconsistent pattern, where asparagine synthesis was missing in LSUCC0396 and LSUCC0226, but present in LSUCC0744 (Figure S3). This likely represents a true asparagine auxotrophy among LSUCC0396 and LSUCC0226, and also indicates that auxotrophies in SAR116 may be strain specific.

DMSP is an important metabolite in SAR116 sulfur metabolism [29, 30]. In this updated analysis, we found DMSP demethylation via *dmdA* encoded across the clade (Figure 4), except for subclade Ib.III. Additional DMSP catabolism genes, *dmdB* and *dmdC* were also encoded across the clade. The *dmdD gene,* encoding an Enoyl-CoA hydratase (ECH) that synthesizes methanethiol and completes the DMSP demethylation pathway, was sparsely distributed across the SAR116 clade (only in subclades Ia.I, Ia.II, and Ib.IV). However, other ECHs such as acryloyl-CoA hydratase (*acuH)* have a broad substrate specificity and have been shown to act on methylthio acryloyl-CoA, the substrate for *dmdD,* and thus *dmdD* and *acuH* are functionally redundant in DMSP catabolism [84–86]. We found that *acuH* genes were more widely dispersed throughout SAR116, occuring in subclades Ia.II, Ia.IV, IbI-III, II and III. Additionally, SAR116 genomes seldomly encoded both *dmdD* and *acuH*, suggesting these act as analogous substitutions for each other. Overall, all subclades except Ib.III were predicted to carry out DMSP demethylation. On the other hand, DMSP lyase (encoded by *dddL, Q, P, D, K,* and/or *W),* a carbon acquisition strategy that releases dimethyl sulfide (DMS), was only detected in subclades I and III and not at all in subclade II. We also found no genetic evidence that SAR116 carry *dmoAB* to convert DMS to methanethiol (Figure 4), therefore it is probable that DMSP cleavage to DMS through DMSP lyase is likely a dead-end for this pathway in SAR116 [87].

**Figure 4:**
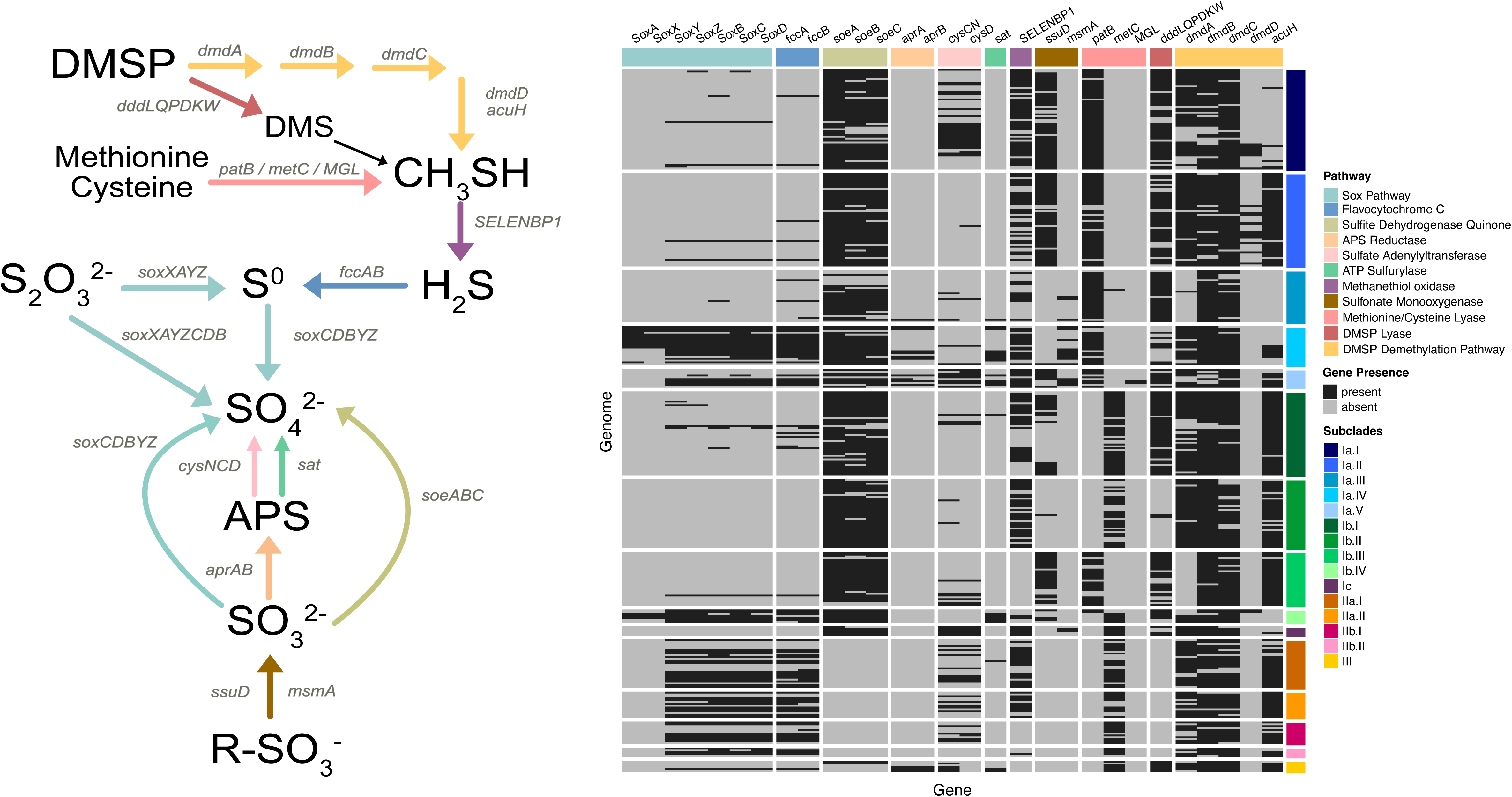
Sulfur oxidation gene heatmap and metabolic pathway map. To the left, the metabolic pathway map, reaction arrow colors match metabolic pathway color represented in the gene presence and absence heatmap to the right. Every row represents a genome, grouped by subclade, and every column represents a sulfur oxidation gene, grouped by pathway. Gray boxes indicate the gene to be absent and black boxes indicate the gene to be present.

SAR116 harbor a variety of genes involved in sulfur oxidation [30, 88]. The potential for thiosulfate oxidation to sulfate via the SOX pathway has been previously highlighted [30], however, in our updated analysis the majority of *SOX*-encoding SAR116 genomes did not carry a complete pathway, specifically, we found no evidence of *soxAX* in any subclade II or III genomes (Figure 4). Only members from subclades Ia.IV and Ib.IV encoded the full pathway (*soxXAYZCDB*), including the complex that binds thiosulfate (*soxAX*) [89]. In subclades II and III, and a few genomes across subclade I, the gene suite contained *soxCDBYZ* that likely oxidizes elemental sulfur or other other sulfane sulfur species to sulfate, and may also oxidize sulfite to sulfate [89]. Thus, not all *SOX*-containing SAR116 genomes have the potential to oxidize thiosulfate. In addition, the SAR116 members that carry the predicted *soxCDBYZ* gene suite, and other members of the SAR116 clade, encode for flavocytochrome C (*fccAB*) that oxidizes hydrogen sulfide to elemental sulfur (Figure 4). Therefore, in these subclades (II and III), hydrogen sulfide may provide the sulfur source that ultimately gets oxidized to sulfate via the *soxCDBYZ* gene suite, which is significant for predicting interactions with sulfur compounds and intracellular sulfur cycling.

We also predicted that some SAR116 can oxidize sulfite to sulfate using the sulfite dehydrogenase quinone, *soeABC* (Figure 4). Aside from two genomes in subclade III, we found this gene suite exclusively in subclade I. Sulfonates that undergo conversion to sulfite via a sulfonate monooxygenase (*ssuD, msmA*), likely supply sulfite to the *soeABC* complex. The *msmA* gene has been previously highlighted [30], however without including *ssuD* an accurate distribution of sulfonate metabolism is lost because subclade I exclusively encodes either SsuD or MsmA (Figure 4). Both proteins act on methanesulfonate, however SsuD can also act on alkanesulfonates [90]. All together, these predicted metabolisms point towards sulfonates as important organosulfur compounds that SAR116 subclade I may be metabolizing to glean additional energy through lithotrophic sulfite oxidation.

The extensive presence of sulfur oxidation genes in SAR116 prompted us to examine their evolutionary history. The key sulfur oxidation genes *soxB* and *fccB* show evidence of vertical transmission throughout the SAR116 clade, whereas *soeA* may have been horizontally transferred extensively throughout subclade diversification (supplemental text).

### Habitats

Metagenomic recruitment of 1,049 marine metagenomes to the 349 SAR116 genomes showed that SAR116 subclades displayed different habitat preferences according to the three marine environments: estuarine, coastal, and open ocean (Figure 5a). Broadly, subclade III represents the dominant group in open ocean systems, followed by subclade II and then subclade I (Figure S5). Spatial distributions at the major subclade level only showed differences in open ocean systems (Figure S5), however when we looked closer at the diversity within subclades I and II, there were clear habitat preferences (Figure 5). Subclades III, IIa.I, and IIb.I dominated in open ocean systems, and the remaining subclades broke into multiple groups with decreasing relative abundance (Figure 5b). In contrast, subclades Ia.IV, Ib.I, and Ia.III were the most abundant groups in coastal systems (Figure 5c). Finally, in estuarine systems, subclades Ia.III and Ia.IV (for which we have three isolates) were the dominant SAR116 subclades (Figure 5d). Thus, habitat diversity in SAR116 mainly occurs at the genus or similar level, which needs to be accounted for to understand how metabolic strategies are linked to preferences for open ocean, coastal, or estuarine systems.

**Figure 5:**
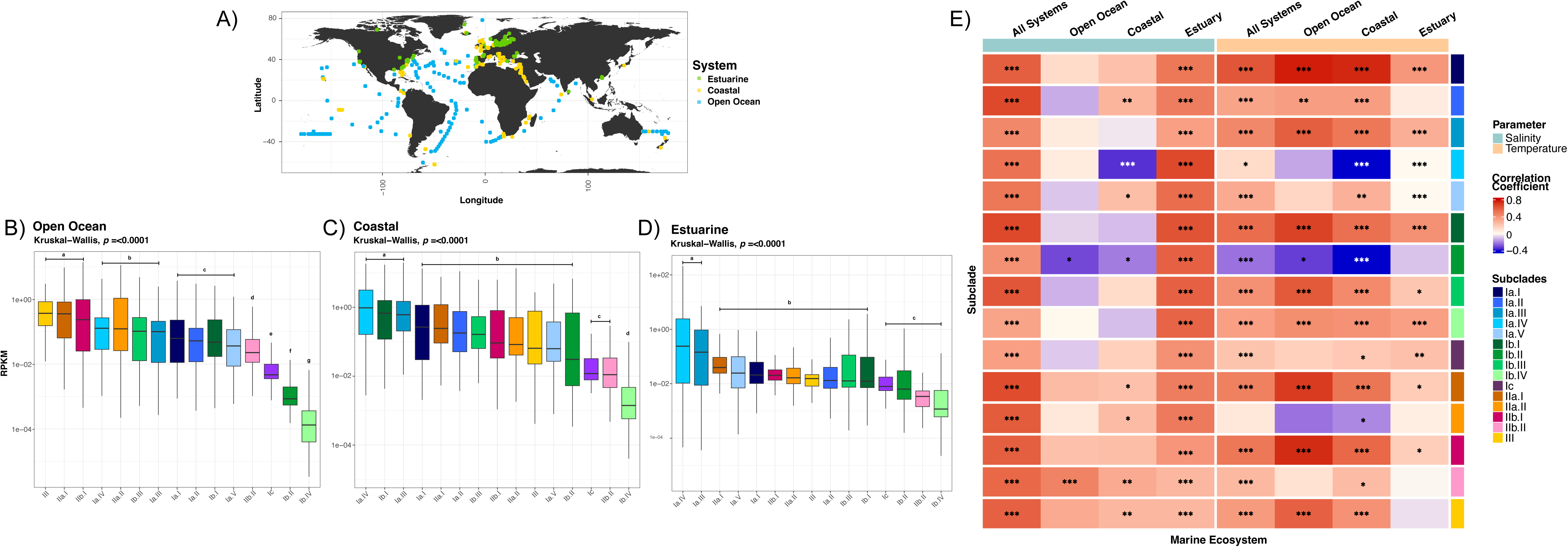
Relative abundance of subclades by ecosystem from metagenomic recruitment of 1,059 metagenomes to all 349 SAR116 genomes. A) World map showing the spatial distributions of the metagenomic samples used for recruitment to the SAR116 genomes. Every data point is a metagenomic sampling site, and marine environments are represented by color. Metagenomes were classified as Open Ocean (B), Coastal (C), or Estuarine (D). Subclades are ordered from the most abundant to least abundant subclade based on median log10-transformed RPKM values. RPKM = reads per kilobase of genome per million mapped reads. Statistical groupings for subclades that share similar mean values are reported above boxplots in panels B-D.

Since we observed specific subclades that were enriched in coastal and estuarine systems where salinity can be a primary differentiator of microbial communities, we compared osmolyte synthesis and transport genes across the SAR116 clade (Figure 3), because the distribution of these genes can provide hypotheses about how prokaryotes mitigate osmotic stress [55, 91, 92]. SAR116 carry genes to synthesize the non-protein amino acid N-`∂-acetyl-ornithine, and sugar derivative di-`*myo*-inositol phosphate, both demonstrated to serve as osmolytes under increasing extracellular salt conditions [93, 94]. The SAR116 clade can also uptake potassium via the *trk* potassium transport system, another common strategy for osmotic stress among prokaryotes [94]. The above pathways are conserved across the SAR116 clade (Figure 3) and represent core osmotic stress strategies for SAR116.

Some osmolyte genes appeared related to habitat differentiation among the subclades. Subclade I, which comprised groups that were more abundant in coastal and estuarine areas, encoded nearly twice as many pathways related to osmotic stress response compared to subclades II and III (Figure 3); genes predicted to be involved in both *de novo* synthesis and uptake of osmolytes. Such an expansion in osmolyte-related genes could indicate that these microorganisms are better adapted to salinity fluctuations, and thus why we find them in greater abundance at coastal and estuarine sites.

Furthermore, subclade I had a unique enrichment of ionic symporters/antiporters (Figure 3). Members of subclade Ib carried a sodium/proton antiporter, and interspersed across subclade I was a sodium/solute symporter. Notably, we found a monovalent cation/H^+^ antiporter exclusively in the genomes of the most abundant subclades in estuarine environments, subclades Ia.III and Ia.IV. Each of these ionic symporters/antiporters have roles in managing osmotic stress [95–97]], and the monovalent cation/H^+^ antiporter has been proposed as conferring an adaptive response to salinity fluctuations in estuaries [28]. Thus, these symporters and antiporters may contribute to the success of subclades Ia.III and Ia.IV in estuarine systems.

Conversely, the few genes encoded by subclades II and III to mitigate osmotic stress are primarily involved in *de novo* synthesis of osmolytes rather than rapid uptake from the environment (Figure 3). This strategy was also seen in *Vibrionaceae* in high salinity when extracellular osmolytes were limited [92], and indicates a potential strategy in the oligotrophic open ocean for subclade II. Specifically, subclade II harbored genes for synthesizing an additional sugar derivative, sorbitol. Subclade III, however, had a balanced representation of synthesis and transport of osmolytes, a strategy seen among other halotolerant microbes [55, 98]. In addition to the aforementioned clade-wide strategies, they also are predicted to synthesize glycine-betaine, along with the transport of ectoine and spermidine/putrescine (Figure 3), potentially serving as additional osmoprotectants or as carbon and nitrogen sources [91, 99]. Overall, the varied strategies to mitigate osmotic stress among SAR116 subclades highlights potential physiological responses and adaptations to environmental stressors throughout SAR116 evolution.

In addition to location, we evaluated the relationships between salinity, temperature, and subclade abundance. Broadly, SAR116 subclade abundances were positively correlated with salinity (Figure 5e, S6), especially in estuarine systems (Figure 5e S7), however salinity did not correlate well with subclade abundances in coastal (Figure S8) or open ocean (Figure S9) environments, potentially due to the small ranges in salinity among these sites. On the other hand, SAR116 subclades appeared more differentially adapted to temperature (Figure 5e, S10). Subclades Ia.I, IIa.I, IIb.I, and III abundances were most positively correlated with temperature, with the strongest correlations in coastal (Figure S11) and open ocean (Figure S12) environments. Subclades Ia.IV, Ib.II, and IIa.II were negatively correlated with temperature (Figure 5e, S10-13), suggesting potentially cold-adapted ecotypes. In estuarine systems, there were no strong correlations between subclade abundances and temperature (Figure S13). Subclades Ia.III, Ia.IV, and Ib.I, the most abundant subclades in coastal systems (Figure 5c), had different relationships to temperature in coastal systems; where subclades Ia.III and Ib.I were positively correlated with increasing temperature, subclade Ia.IV was negatively correlated with increasing temperature (Figure S11), suggesting that subclades Ia.III and Ib.I are the dominant SAR116 coastal subclades in warmer regions, and subclade Ia.IV in cooler regions. Collectively, this indicated that temperature may drive SAR116 spatial distributions and habitat preferences.

### Isolate physiology and morphology

We characterized key growth parameters for three isolates representing the breadth of cultured SAR116 diversity: LSUCC0719 (Ia.I), LSUCC0744 (Ia.IV), and LSUCC0684 (Ic). All could grow in salinities ranging from 11.6 - 34.7 ppt and had no discernable differences in growth rate across this range, with optimum growth occurring at 23.2 ppt (Figures 6a, S14). However, each isolate had a specific temperature tolerance range and optimum temperature for growth (Figures 6b,c; S15). These isolates all had maximum growth rates under the conditions tested between roughly one and two doublings per day (Figure 6c).

**Figure 6:**
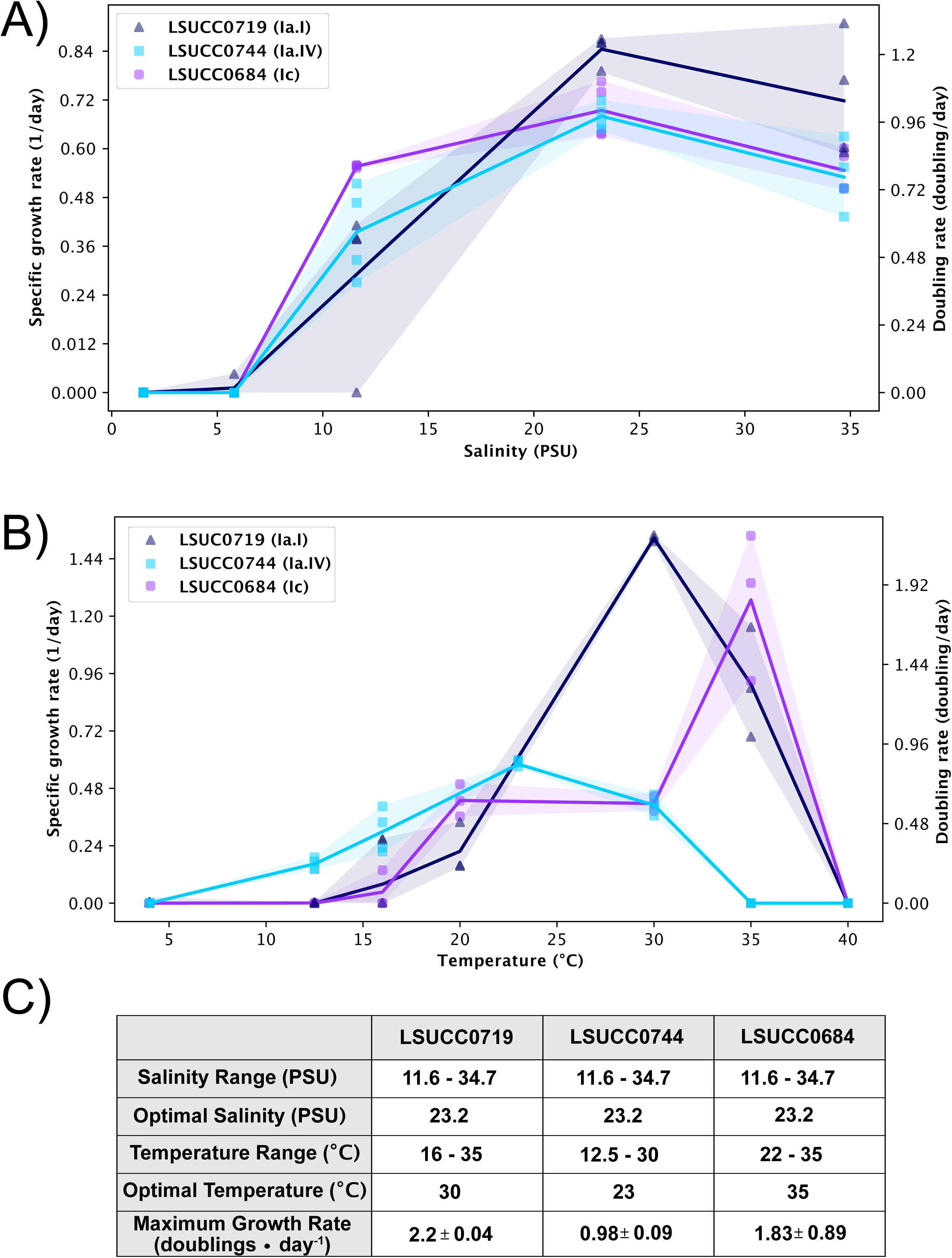
Phenotypic variation of SAR116 LSUCC isolate representatives across temperature and salinity ranges. LSUCC0719, representing subclade Ia.I in dark blue. LSUCC0744, representing subclade Ia.IV in light blue. LSUCC0684, representing subclade Ic in purple. A) Salinity vs. Growth rate. B) Temperature vs. Growth rate

**Figure 7:**
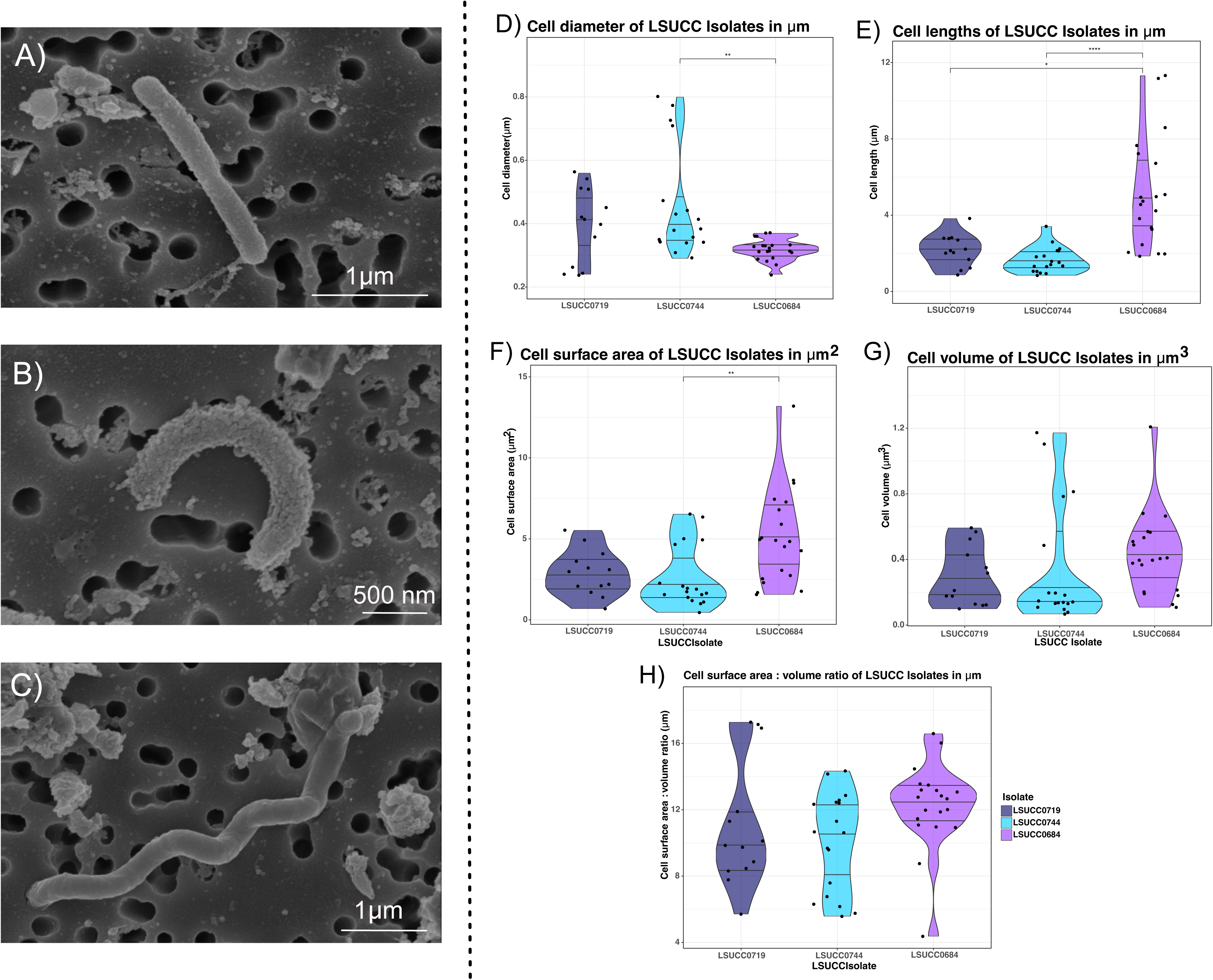
Cell sizes and morphologies for SAR116 isolates, LSUCC0719 (n=13 cells), LSUCC0744 (n=18 cells), LSUCC0684 (n=20). Measured cell diameters (A), cell lengths (B), cell surface areas (C), cell volumes (D), and cell surface area:volume ratios. Representative SEM images for LSUCC0719 (F, scale bar = 1*μ*m), LSUCC0744 (G, scale bar = 500nm), LSUCC0684 (H, scale bar = 1*μ*m). ‘*’ = p-value < 0.05, ‘**’ = p-value < 0.01, ‘****’ = p-value < 0.0001.

Our SAR116 isolates each displayed different morphologies, covering bacilli, (LSUCC0719, Figure S16), vibrio (LSUCC0744, Figure S17), and spirilla (LSUCC0684, FigureS18) (Figure 6f-h). Although their morphologies were different, LSUCC0719 and LSUCC0744 showed no significant differences in their cell dimensions (Figure 6a-e), while LSUCC0744 and LSUCC0684 had differences in cell diameter (Figure 6a) and surface area (Figure 6c). The greatest variation in cell dimensions occurred across cell lengths (Figure 6b), where LSUCC0684 had a considerable range between 1.85 - 11.32*μ*m (Figure S2). Nevertheless, despite the variation in morphologies and cell sizes, there were no significant differences in cell volume (Figure 6d), or cell surface area to volume ratios (Figure 6e) between these cells, suggesting a conservation of these characteristics.

We also cataloged extensive pleomorphy that may be related to a complex life cycle in the host-associated (Ic) representative, LSUCC0684 (supplemental text). These cells appeared both as long spirilla, but also as flattened large cocci (Figure S22b-d). We also captured an image of LSUCC0684 undergoing a considerable morphological change from the spirillum to the cocci form (Figure S22a). Thus, we believe these organisms are pleomorphic and the image of the cell spanning both morphologies may be a visualization of the conversion to the endosymbiotic calcifying SAR116 cells found in marine sponges [82], termed “calcibacteria”. These endosymbiotic “calcibacteria” SAR116 cells were more spherical in shape, approximately 1*μ*m in diameter, in sponge tissues [82]. This matches our measured diameters of the spherical LSUCC0684 morphology ranging from 0.72 - 1.25*μ*m in diameter (Figure S22b-d). Some appeared to be hollowed or collapsed (Figure S22b). We hypothesize that this morphological transformation occurs as LSUCC0684 cells enter stationary phase, which may correspond to a life cycle change where they are taken up by sponge hosts [82].

## Discussion

Our isolation of new, diverse SAR116 strains motivated us to investigate the *Puniceispirillales* in the context of new genomes and the first comparative phenotypic data for the group. In addition to five new strains in three putative genera, we contributed the first four closed SAR116 genomes and used them to anchor the largest SAR116 pangenomic study to date. These closed genomes allowed us to make more confident predictions of auxotrophy, and our study updated SAR116 phylogeny, provided an updated description of metabolic heterogeneity within the clade, and also the first evidence for subclade-specific environmental diversification.

We found that a notable difference in SAR116 subclades was temperature differentiation. We observed this both within environmental abundances, via metagenomic recruitment (Figure 5e), and corroborated some of these patterns through experimentation with our isolates (Figure 6b). Subclade Ia.I positively correlated with temperature across all marine systems, particularly in open ocean and coastal systems, and the representative isolate strain LSUCC0719 also had a temperature growth optimum of 30°C. Similarly, our warmest-adapted isolate, LSUCC0684 (temperature growth optimum of 35°C), belonged to subclade Ic, which were also positively correlated with temperature in coastal and estuarine systems. This group is also the subclade predicted to have a host-associated life cycle, and thus the 35°C temperature optimum aligns with endosymbiotic relationships with corals and sponges [66, 82] where host temperature tolerances can range 23-32°C [100]. Conversely, our more cold-adapted strain, LSUCC0744, belonged to subclade Ia.IV, which was negatively correlated with temperature. The temperature range for LSUCC0744 also matched that of the previously isolated Ia.IV strain IMCC1322 (12.8 - 34.2^ο^C) [13]. Maximum growth rates of the LSUCC SAR116 cultured representatives indicate they fall in the spectrum between the canonical slow growth of SAR11 [55], and faster growers such as members of the Oligotrophic Marine Gammaproteobacteria (OMG) [101]. Without cultured representatives from subclades II and III, we do not definitively know their growth temperature ranges, however our data suggested that they are positively correlated with increasing temperature, particularly the members of subclades II and III. While these open-ocean groups may be resilient to increasing sea-surface temperatures, the dominant coastal and estuarine subclade Ia.IV that prefers cooler temperatures, may face displacement. Thus, SAR116 contributions to coastal and estuarine biogeochemistry may be altered in a changing climate, for example, subclade Ia.IV was one of the only groups predicted to carry out thiosulfate oxidation.

Isolate physiology also supported the assertion that the dominant coastal and estuarine subclade Ia.IV was adapted to lower salinities. Strain LSUCC0744 had an optimum growth salinity of 23.2 ppt, which corroborates previous reports from the other Ia.IV isolate strain IMCC1322, that had an optimum of 28 [13] Our other isolates, all belonging to the larger subclade I, also had the same optimum salinity (Figure 6b). Subclade I comprised groups that dominated in coastal and estuarine systems, so these salinity optima corroborate those patterns. We now know from our distribution data (Figure 5) that the coastal sampling locations yielding the LSUCC, HIMB, and IMCC isolates also meant that obtaining subclade I taxa was more likely than those from subclades II or III [7, 13, 18, 76]. This predicted (Figure 5e) and observed (Figure 6b) preference for brackish waters in subclade I may result from the wide array of osmotic stress response systems encoded by subclade I genomes (Figure 3), that likely improve tolerance to salinity fluctuations in coastal and estuarine systems [102], particularly through rapid uptake of positive and negative ions [92].

We can use our new characterizations of habitats and metabolic preferences to contextualize previous ecological observations of SAR116. These organisms were most abundant in the Columbia River estuary system among brackish salinities between 15.4 and 25.4, and showed high expression of metallopeptidases, along with zinc, iron, and nickel transporters [28]. Phenotypic responses (Figure 6b), salinity correlations (Figure 5e), and habitat preferences (Figure 5c&d) suggest that these metabolically active SAR116 members were from subclade I. Our pangenomics results can even help us define them further as members of subclade Ia.IV or Ia.V since these Ia groups carried zinc, nickel, and iron transporters (Figure 3). Separately, SAR116 were the most abundant *dddP*-carrying taxa in a northwest Pacific transect, separated into two *dddP* OTUs with differing biogeographical patterns: OTU43 was abundant in both coastal and open-ocean sites and OTU28 was only detected in the open-ocean [29]. We found no evidence for *dddP* in subclade II (Figure 3), and given the preference of subclade III for open-ocean systems (Figure 5b) compared to the coastal/open-ocean subclade I habitats (Figure 5b&c), we predicted OTU43 to be a member of subclade I and OTU28 to be a member of subclade III. Indeed, a blastn of the *dddP* sequences from that study against the LSUCC0719 and AG_899_J15 genomes, representing subclades I and III, respectively, showed that our subclade I representative had 545 dddP sequence hits, with 19 sequences greater than 90% identical, whereas the subclade III representative had 44 *dddP* sequences hits with only one sequence greater than 90% identical. Thus, subclade I likely dominated the *dddP*-containing SAR1116 population in this region.

DMSP lyase is one of several types of sulfur metabolism that were differentially distributed across the SAR116 clade (Figure 4). Different groups can access a variety of sulfur compounds which likely links them to phytoplankton in distinct ways. Sulfonates are produced by numerous phytoplankton groups in coastal and open ocean systems [103]. Sulfonate monooxygenase (*ssuD, msmA*) genes that convert sulfonates to sulfite were encoded exclusively by subclade I (Figure 4), suggesting that sulfonates are a phytoplankton-supplied source of sulfite for these organisms. Additionally, while most SAR116 have the genetic capability to demethylate DMSP, only subclades I and IIa show the genetic capability to oxidize hydrogen sulfide derived from DMSP via methanethiol oxidase (Figure 4). Thus, we hypothesize subclades I and IIa interact with both the organic and inorganic sulfur pools via multiple organic sulfur species likely originating from phytoplankton. For subclades that encode flavocytochrome C (*fccAB*) that produces elemental sulfur from hydrogen sulfide (Figure 4), if that elemental sulfur is not immediately oxidized to sulfate (*soxBCDYZ)*, then it may accumulate intracellularly as sulfur inclusions for later oxidation. This may confer an advantage to these SAR116 subpopulations, particularly in the oligotrophic open oceans, and potentially operate as a sulfur-sink in the surface ocean.

A critical component to determining the relative contributions of microbial cells to biogeochemical cycles is their sizes, and this work provides a new, more comprehensive view of morphological diversity and cell dimensions for SAR116. Previous data showed considerable variation in cell morphology across two different isolates, IMCC1322 (Ia.IV) [13] and HIMB100 (Ib.I), suggesting that SAR116 may display a wide range of cell shapes and sizes. We corroborate the vibrioid shape of subclade Ia.IV with LSUCC0744 (Figure 6b), and both isolates have similar cellular dimensions. Strain HIMB100 (Ib.I) had a spirillum morphology with cells ranging 1 - 5*μ*m in length [18], more similar to that of LSUCC0684 (Figure 6c), representing subclade Ic (Figure S4). Conversely, strain LSUCC0719 had a more simple bacillus shape compared to the other cell types. Thus, within just subclade I, there were at least three distinct morphologies across a range of sizes. LSUCC0719 and LSUCC0744 representatives were large relative to other cultivated marine microbes like members of SAR11 [43], OM252 [42], but were more similar in size to members of the *Roseobacter* lineage [104]. The conserved surface area:volume ratio despite varied sizes (particularly cell length, Figure 6e) and morphologies suggests that SAR116 might regulate cell size relative to their surface area:volume ratio, and/or this was a maintained characteristic shared by their common ancestor because of its importance for nutrient acquisition [92].

The range in cell length for LSUCC0684 was large (Figure 6e, Figure S2), and may therefore represent a strain capable of filamentous growth, similarly to sulfide-oxidizing *Beggiatoa spp.* [105]. LSUCC0684 displayed significant pleomorphy, so we hypothesize that filamentous growth may occur in preparation for the next stage in the LSUCC0684 life cycle where cells increase in size prior to undergoing a morphological transformation (Figure S18), potentially in anticipation for uptake by corals and sponges [66, 82]. Endosymbiotic *Proteobacteria* in marine sponges aid in larval settlement, a process regulated by nitric oxide signaling [106] via arginine biosynthesis that stimulates the sponge host to produce nitric oxide and thus initial larval settlement [107]. Subclade Ic were prototrophic for arginine (along with all other amino acids- Figure 3), and therefore amino acid production may be a role for endosymbiotic SAR116 Ic cells in marine sponges. Further work should explore the involvement and activities of SAR116 Ic in sponge and coral holobionts. Other endosymbiotic lineages, such as squid endosymbiont *Vibrio fischeri* and jellyfish endosymbiont *Symbiodinium microadriaticum* [108], each have planktonic phase of their life-cycle [109, 110]. Therefore, at least transient endosymbiotic relationships of what we consider “free-living” organisms may be occurring more frequently than previously thought.

Altogether, this investigation demonstrated SAR116 diversification through metabolic and spatial niche partitioning, anchored by information from new and diverse SAR116 isolates. Our experimental results expanded the known salinity and temperature tolerances across subclade I and corroborate culture-independent assessments of their habitat preferences and spatial distributions. Our cultures also revealed large differences in cell dimensions and shapes throughout subclade I, and identified a consistency of surface area:volume ratios despite large differences in cell sizes. We were also able to identify a peculiar morphological change in the subclade Ic representative, LSUCC0684, that may link these organisms to a host-associated part of their life cycle. This isolate provides new opportunities to investigate symbioses of SAR116 with animals, a very understudied aspect of SAR116 biology. The diversity of SAR116 leaves open many other avenues for investigation: future research to validate the hypothesized sulfur oxidation metabolisms, osmotic stress regulation, and putative vitamin and amino acid auxotrophies will be important to deepen our understanding of how SAR116 physiology influences sulfur cycling, interacts with other members of the microbial community, and differentiates subclades in relationship to salinity variation.

## Data Availability

Assembled isolate genomes for LSUCC0226, LSUCC0396, LSUCC0684, LSUCC0719, and LSUCC0744 are available on NCBI under CP166132, CP166131, CP166130, JBFPJN000000000, and CP166129, respectively. Raw reads are available under BioProject PRJNA1133775. The accessory data sheets including the pangenome summary are available at: https://figshare.com/account/home#/projects/233330. Cryostocks of isolates used in this analysis are available upon request.

## Acknowledgements

We would like to thank the former USC Genome Core for Illumina NextSeq sequencing of LSUCC isolate genomes. We would like to specifically thank Suchi Patel for performing library preparation and sequencing. We would also like to thank Dr. Ben Temperton at the University of Exeter for base-calling and quality checking our MinION long read sequences. The authors would also like to thank the University of Southern California (USC) Core Center of Excellence in Nano Imagining (CNI) for the availability of scanning electron microscopes, and Carolyn Marks for sample preparation and imaging. We would also like to acknowledge the Center for Advanced Research Computing (CARC) at the University of Southern California (https://carc.usc.edu) for computational resources that have contributed to the results in this publication. This work was supported by a Simons Early Career Investigator in Marine Microbial Ecology and Evolution Award, and NSF Biological Oceanography Program OCE-1945279 and Emerging Frontiers Program EF-2125191 grants to J.C.T., and a Lerner Gray grant from the American Natural History Museum awarded to M.W.H.

## Contributions

JTC conducted the analysis, experimentation, generated figures and wrote the paper. LT, MWH, VCL, CYK contributed strains, data, or assisted in experimentation. JCT devised the study, obtained funding, and assisted in manuscript preparation. All authors contributed to edits in the final manuscript.

