## Supplemental text and figures for "Culture-supported ecophysiology of the SAR116 clade demonstrates metabolic and spatial niche partitioning"

### Methods

*Isolation, genome sequencing, and assembly*

We isolated five SAR116 strains- - LSUCC0226, LSUCC0396, LSUCC0719, LSUCC0744, and LSUCC0684- using high-throughput dilution-to-extinction cultivation and identified them via 16S rRNA gene comparisons, as previously described [1]. LSUCC0226 was isolated on October 1, 2015 from the Calcasieu Jetties in Cameron, LA (29.760164 -93.340159), and LSUCC0396 on April 6, 2016 off of Freshwater City Road in Vermilion Parish, LA (29.53084, -92.326147), from the both using JW2 media [1]. LSUCC0684 and LSUCC0719 were isolated on August 22, 2016 from the Calcasieu Jetties in Cameron, LA (29.760164 -93.340159), and LSUCC0744 was isolated on November 11, 2016 off of Freshwater City Road in Vermilion Parish, LA (29.53084, -92.326147) each using MWH2 media [2]. For genome sequencing, we grew these isolates axenically in a range of 500mL – 1.5 liters of MWH2 medium from cryostocks [2]. Cells were filtered onto 0.2 μm polyethersulfone (PES) filters (Pall Corporation, New York, USA) and stored at -20˚C for genomic DNA extraction. We extracted genomic DNA using the GenElute Bacterial Genomic DNA Kit (Sigma-Aldrich, Missouri, USA), then cleaned and concentrated the DNA with the Zymo Genomic DNA clean and concentrator kit. Illumina NextSeq 550 library preparation was done via a KAPA HyperPrep kit, and sequencing was performed at the University of Southern California (USC) Molecular Genomics Core. To obtain additional DNA for Oxford Nanopore sequencing, we grew separate cultures of LSUCC0226, LSUCC0396, LSUCC744, and LSUCC0684 from cryostocks in 1 L MWH2. DNA extraction, cleaning, concentrating, and quantification was done as described above. DNA was sheared to a 20 kbp fragment size using Covaris g-TUBE (Covaris, Inc. Massachusetts, USA). Quality assessment of the sheared DNA was done at the USC Molecular Genomics core using the Agilent 4200 TapeStation. Library preparation began with 700 ng of DNA per isolate strain and was performed using a modified 1D Native barcoding genomic DNA (with EXP-NBD103 and SQK-LSK108) protocol. Prior to sequencing, we checked the pore activity of the R9.4.1 FLOMIN106 flow cell and configured it using the MinKNOW 3.1.19 software. We loaded 300 ng of pooled DNA onto the flow cell, and sequenced with a MinION for 20 hours generating 174.26 k reads and 811.18 Mbp. Prior to assembly, Nanopore reads were base-called using Guppy v330 (Oxford-Nanopore Technologies) with GPU configuration to increase base-calling speed, and a minimum quality score of 7.0. We demultiplexed Nanopore reads using Porechop [3] under the stringent binning settings, with the additional option to remove barcode adapters in the middle of the read. Illumina reads were first trimmed using Trimmomatic v0.38 [4] to remove bases less than 20 on a Phred-33 scale. Leading and trailing bases of a read were removed if below the threshold of 20, and bases were cut if the average quality score fell below 20 on a sliding window of 13 bases. Additionally, a read was removed if it was below 40 bp in length. We hybrid-assembled genomes that received both Illumina and Nanopore sequencing using Unicycler v0.4.8 [5] on default settings. The genome of LSUCC0719 only received Illumina sequencing and we assembled the trimmed reads using SPAdes v3.13.1 [6] with the automatic coverage cutoff specified. We assessed genome assembly quality and completeness using CheckM v.1.1.3 [7] using lineage_wf settings for both hybrid and Illumina-only assemblies.

*Genomic Relatedness*

We used average nucleotide identity (ANI) and average amino acid identity (AAI) to delineate SAR116 subclade structure and to assess the relatedness of subclade members. We computed ANI across SAR116 members using FastANI [8], and computed pairwise AAI across SAR116 members using CompareM [9], both with default settings. To determine the divergence of the overall clade, the 16S rRNA gene sequences from the two most divergent genomes were compared via BLAST [10]. We evaluated the relationship between 16S rRNA gene identity, phylogenomic tree topology, and AAI to confirm clade structure and ultimately determined subclade delineations with AAI [11][12].

*Environmental preferences*

To understand the relationship between SAR116 subclades and major marine environments we performed metagenomic read recruitment using as previously described [13, 14], using Reads-Per-Kilobase-Mapped (RPKM) as a proxy for subclade abundance. RPKM values were calculated for each individual genome using RRAP [15], and then RPKMs for all the genomes in a subclade were summed to obtain subclade RPKM. We classified the sample sites across the 1,059 metagenomes [14] as Open Ocean, Coastal, or Estuarine. A metagenome was considered “Coastal” if it was collected from within the distance of the local coastal shelf and considered “Estuarine” if it was collected within an estuary. All other metagenomes were considered to be “Open Ocean”. Salinity from sites associated with metagenomic data was reported as unitless, as these measurements arise from a conductivity ratio. We used a Kruskal-Wallis test [16] to determine if there were statistically significant differences between SAR116 subclade abundances and salinity of the metagenomic samples. To quantify pairwise subclade abundance differed based on salinity values, a Dunn’s test [17] was employed. Subclade abundance data based on marine environment was visualized using custom R-scripts made in R-Studio [18, 19].

*Single Gene Phylogenetics*

We compared our updated SAR116 taxon selection to previously described SAR116 16S rRNA gene subclade structure using 16S rRNA genes extracted from the Anvi'o database via 'anvi-get-sequences-for-hmm-hits' and sequences from the 16S rRNA gene phylogenetic tree in Lee et al. 2019 [20]. To investigate SAR116 association with corals and sponges, we additionally obtained 16S rRNA genes in Bonthond et al. 2018 [21] from the SILVA database [22–24]. Sequences were trimmed with TrimAl [25] and aligned with MUSCLE [26], both with default settings. Maximum-likelihood inference was done using IQ-TREE v2.1.2 [27] using the automated nucleotic acid substitution best-fit model estimator '-m MFP' and traditional bootstrapping with 1000 replicates. The selected nucleic acid substitution model was TVMe+R5, and the resulting tree was visualized using *Ggtree* v3.2.1 and *Treeio* v1.18.1 R packages [28–30], rooted at the midpoint, and nodes were ordered in increasing order.

We used SoxB within the thiosulfate oxidizing SOX pathway for phylogenetic inference as it is the gene responsible for releasing sulfate at the end of the pathway, and is the marker gene for full thiosulfate oxidation [31]. The active site encoding subunit, f*ccB* in the flavocytochrome c (*fccAB)* complex [31], and the active site encoding subunit, *soeA*, within the sulfite dehydrogenase quinone (*soeABC)* complex [32] were also chosen for the phylogenetic inference of sulfide oxidation and sulfite oxidation, respectively. Amino acid sequences for all genomes were generated in Anvi'o v7.1 using 'anvi-get-sequences-for-gene-calls'. For each sulfur oxidizing gene, the corresponding amino acid sequences were searched against the NCBI RefSeq database [33] via BLAST [10]. The top 25 best-hit were retained for each query, and then further filtered to remove any duplicates. This resulted in 166 SoxB, 313 FccB, and 672 SoeA amino acid sequences for phylogenetic inference of sulfur oxidizing genes. Sequences were aligned, trimmed, and phylogenetic inference and visualization was completed as above. The amino acid substitution models were LG+I+G4, LG+G+R6, LG+R9 for SoxB, FccB, and SoeA, respectively.

*Temperature, and Salinity Tolerances*

We tested the growth ranges and optima for temperature and salinity using one isolate from each of the three phylogenetic groups for which we had cultures. Although we have five SAR116 isolates, LSUCC0744 was chosen as the sole representative of IIa.I since it shares > 98% 16S rRNA gene identity with LSUCC0226 and LSUCC0396. To assess the temperature growth ranges across these isolates, each isolate representative was inoculated from late-log phase in triplicate into 5 mL of MWH2 in 10mL sterile borosilicate glass test-tubes (#1512, Globe Scientific, New Jersey, USA) and incubated in the dark at 4, 12, 16, 22, 30, 35, and 40$^{o}$C. Salinity growth ranges were also tested across these isolates, and similarly, each isolate representative was inoculated from late-log phase in triplicate into 5 mL of modified MWH artificial seawater medium with salinities ranging from 1.5 - 34.7 ppt (MWH1 - MWH5) into 10mL sterile borosilicate glass test-tubes, and incubated in the dark at 23$^{o}$C. Salinity for the growth experiments was calculated based on the chlorinity (salinity (ppt) = 1.80655 x Cl (ppt) [34] of the "base salts". Growth for all experiments was measured with the BD Accuri C6 Plus flow cytometer (Becton, Dickinson and Company, New Jersey, USA) using 1x SYBR Green as described [1, 13] and growth rates were calculated using sparse-growth-curve [35]. To contextualize these phenotypic responses to salinity and temperature, we compared the metagenomic recruitment to the summed RPKMs for each isolate clade based on salinity and temperature across all metagenomic datasets. We fit linear regressions based on salinity or temperature vs. subclade log-transformed RPKM and visualized the results using 'ggplot2' [36].

*Scanning electron microscopy*

Three LSUCC SAR116 isolates – LSUCC0719, LSUCC0744, and LSUCC0684 -were inoculated from cryostocks (1 mL) in 100 mL of MWH2 media [2] in sterile 125 mL flasks (FPC0125S, TriForest, Irvine, CA), and fixed at a final concentration of 2.5% (v/v) glutaraldehyde (G5882, Sigma-Aldrich) after reaching exponential growth. Scanning electron microscopy (SEM) was performed at the USC NanoImaging Center. SEM preparation was performed as described in [14]. Briefly, we filtered cells through a 0.2 micron PES filter (Pall Corporation, New York, USA) and adhered cells to the filter membrane via a poly-L-lysine coating. Cell membranes were stained with 0.1 M HEPES buffered 0.05% Ruthenium Red (RuRed) and 10% sucrose and incubated with cells for 10 minutes. The stain was fixed into cells by a 0.1 M HEPES buffered 0.05% RuRed, 0.8% Osmium tetroxide, and 10% sucrose solution and incubated for 25 minutes. After, we washed the filter membrane with 0.1 M HEPES and 10% sucrose, followed by 10 mL of deionized water. We then dehydrated the filters sequentially at 50%, 70%, 95% and 100% ethanol, then stored in 100% ethanol until imaging.The resulting SEM images were analyzed in ImageJ [37] to quantify cell size measurements.

Cell volume and cell surface area measurements were modeled based on the specific morphology of the cell. Bacillus morphologies, due to their resemblance to a capsule, were modeled using Equations 1 & 2, which are equations for modeling volume and surface area of a capsule (Figure S1). Vibrio morphologies were modeled using Equations 3 & 4 for curved rod shaped cells, or 5 & 6 for horseshoe shaped cells. Curved rod shaped cells were modeled to emulate two hemispheres (Figure S16), and horseshoe shaped cells were modeled to emulate one cylinder with two hemispheres at either end (Figure S17). Spirillum morphologies were modeled using Equations (7 & 8), where every helix was modeled as a cylinder and each end of the cell was modeled as a hemisphere, and overall surface area and volume were a sum of the number of helices plus the cell ends (Figure S18). In Equations 1-8, the variable *h* represents the length of the capsule minus the radius to in order to capture the dome volume and surface area at the end of a capsule. In equations 7 & 8 the variable *n* represents the variable total number of helices per cell.

Equation (1): Volume, Bacillus

$$\pi r^2 (\frac{4}{3} r) + h$$

Equation (2): Surface area, Bacillus

$$2\pi r(2r + h)$$

Equation (3): Volume, Vibrio (curved rod)

$$(\frac{2}{3}\pi r^3)x2 + (\pi r^2 h)x2$$

Equation (4): Surface area, Vibrio (curved rod)

$$(2\pi r^2)x2 + (2\pi rh)x2$$

Equation (5): Volume, Vibrio (horseshoe)

$$(\frac{2}{3}\pi r^3)x2 + \pi r^2 h_1 + \pi r^2 h_2 + \pi r^2 h_3$$

Equation (6): Surface area, Vibrio (horseshoe)

$$(2\pi r^2)x2 + 2\pi rh_1 + 2\pi rh_2 + 2\pi rh_3$$

Equation (7): Volume, Spirillum

$$(\frac{2}{3}\pi r^3)x2 + \pi r^2 h_1 + \pi r^2 h_2 + \pi r^2 h_3 + \ldots \pi r^2 h_n$$

Equation (8): Surface area, Spirillum

$$(2\pi r^2)x2 + 2\pi rh_1 + 2\pi rh_2 + 2\pi rh_3 + \ldots 2\pi rh_n$$

We tested statistical differences in cell measurements by first employing a Bartlett test [38] to check for equal variance in the cell measurements. If variance was unequal between cell measurements, then a Welch's ANOVA [39] was performed to determine if the difference between the mean cell measurements was statistically significant. If variance was equal, then a standard ANOVA was performed [40]. Finally, we used a Dunn's test [17] to determine pairwise differences between cell measurement means. Cell measurements and statistics were visualized using 'ggplot2' [36] and 'rstatix' [41].

### Results

*New metabolic predictions*

In addition to an aa3-type cytochrome-c oxidoreductase, two isolates, LSUCC0744 (Ia.IV) and LSUCC0684 (Ic), also had predicted high-oxygen-affinity oxidoreductases (cytochrome c oxidase: cbb3-type and cytochrome bd complex, Table S5), though these genes were not found among other subclade Ia.IV or Ic representatives, which either indicates that our isolates are unique compared to their close relatives, or that we were able to detect these genes only because we had complete genomes. Members of subclade Ia.III only encoded subunit II of the cytochrome bd complex, and while subunit I can function without subunit II [68], it is unclear if the inverse is true. Alternatively, a novel protein may serve as an analogous substitution for subunit I. While the overall SAR116 clade abundance typically declines with decreasing oxygen [69], the relationship between subclade Ia.III and low oxygen has yet to be established. Regardless, the presence of these high-affinity oxioreductases indicates the potential for SAR116 aerobic metabolism under low oxygen conditions.

Methanethiol oxidase, which transforms methanethiol to hydrogen sulfide, is another gene predicted to supply organic sulfur substrates for downstream inorganic sulfur oxidation (Figure 4). Aside from DMSP (discussed above), methanethiol may also come from methionine or cysteine via the *patB/metC* cysteine ligase, or *MGL* methionine ligase, both of which have shown to form thiols [78], and are involved in methionine salvage pathways [79]. SAR116 genomes almost exclusively carry either *patB, metC,* or *MGL*, suggesting an analogous metabolic function among SAR116 subclades. Aside from subclades Ib.III, IIb.II, and III, methanethiol oxidases are present throughout the rest of the SAR116 subclades (Figure 4). Subclades that do not harbor *fccAB* to oxidize hydrogen sulfide to elemental sulfur likely utilize hydrogen sulfide for cysteine and methionine biosynthesis [80].

*Evolutionary history of sulfur oxidizing genes in SAR116*

SAR116 formed a monophyletic group within the SoxB phylogenetic tree (Figure S19) that was sister to the *Rhizobiales* and *Rhodospirillales*, similarly to the 16S rRNA gene and phylogenomic trees (Figure 1a), suggesting a vertical inheritance of these genes. An unclassified *Rhodobacteraceae* sequence branched within the SAR116 clade and was most closely related to SAR116 IIb.I, indicating a potential misclassification, or horizontal gene transfer. Subclades I and II formed distinct clades, and their branching structures did not exactly match that of the updated phylogenomic tree (Figure S1), implying horizontal transfer from the *Rhizobiales* to SAR116 subclade Ia.IV, that then diversified with SAR116 evolution. Many of the outgroup SoxB sequences belonged to known sulfur oxidizers [42–44], providing stronger evidence for the metabolic potential of thiosulfate oxidation within SAR116.

Similarly, SAR116 formed a monophyletic group within the phylogenetic tree of FccB (which oxidizes hydrogen sulfide to elemental sulfur) (Figure S20). SAR116 are sister taxa to *Rhizobiales,* many of which are known sulfur oxidizers [45], similar to many other prokaryotic taxa in the *fccB* phylogenetic tree [31, 46], which validates the metabolic potential of hydrogen sulfide oxidation with SAR116. Similar to the SoxB phylogeny, the monophyletic structure and branching patterns suggest that *fccB* was horizontally transferred from the *Rhizobiales* to members of subclade I, then further diversified throughout SAR116 evolution

Unlike the SoxB and FccB phylogenies, the SAR116 SoeA (for oxidation of sulfite to sulfate) phylogeny had a polyphyletic structure (Figure S21). Group 1 in the tree, comprised of members from subclades Ia, Ib, and III, were sister taxa to the *Rhodospirillales*; and Group 2, comprised of subclades Ib.IV, Ic, and a member from subclade Ia.IV, were sister to members of the *Rhodobacterales* and *Rhizobiales*. The observed paraphyletic structure indicates that this metabolism has had a different evolutionary path for subclades Ib.IV and Ic than subclades I and III. Additionally, the branching patterns within Groups 1 and 2 are different from the phylogenomic tree (Figure S1), indicating a horizontal gene transfer from the *Rhodospirillales* in Group 1 and the *Rhizobiales* in Group 2. Both members of the SAR116 sister taxa are known sulfur oxidizers which validates this metabolic potential in SAR116.

### Figure Captions

**Figure S1: Phylogenomic tree of the SAR116 clade with other members of the *Alphaproteobacteria* as an outgroup (left), and pairwise Average Nucleotide Identity (ANI) vs. Average Amino acid Identity (AAI) of the SAR116 clade (right).** The scale bar on the phylogenomic tree represents 0.1 changes per position, and bootstrap support values (n=1000) are indicated at nodes. Shapes overlaying internal nodes indicate taxonomic classification where squares represent Family, triangles represent Genus, and circles represent Species. Starts next to tip labels represent isolates based and the color of the star indicates the culture collection they are housed in - red (LSUCC), blue (IMCC), and green (HIMB). Dashed lines delineate subclades within the SAR116 clade and their corresponding ANI v. AAI values. Red circles represent genomes that received pairwise 16S alignment, and pairwise AAI comparison to validate the Order level divergence of the SAR116 clade.

**Figure S2: 16S rRNA gene phylogenetic tree of SAR116 members and outgroup members of the *Alphaproteobacteria*.** The scale bar represents 0.3 changes per position. Bootstrap values (n=1000) are represented at nodes. Black tip labels indicate SAR116 members, red tip labels indicate SAR116 members from water sampled at the coral:water interface, orange tip labels indicate SAR116 members from ground coral tissue samples, and gray tip labels indicate outgroup members. Subclades are labeled with the updated subclade designations from this investigation (left) and also the historical 16S subclade designations (right).

**Figure S3: KEGG-Decoder output heatmap for metabolic pathway completion.** Every row genome that corresponds to phylogenetic ordering based upon the phylogenomic tree in Figure 1. Every column is a metabolic pathway. The darker the color the more complete the pathway, with the pale yellow color indicating no detection and dark red indicating a complete pathway.

**Figure S4: Estimated genome completeness vs. subclade.** Estimated percent completion (percent) of SAR116 assembled genomes by subclade. Estimated genome completeness data were collected from CheckM output.

**Figure S5: Relative abundance of major SAR116 subclade classification by ecosystem from metagenomic recruitment of 1,059 metagenomes to all 349 SAR116 genomes**. Broad subclade classifications are ordered from the most abundant to least abundant subclade based on mean log10-transformed RPKM values. RPKM = reads per kilobase of genome per million mapped reads. Metagenomes were classified as Estuarine (A), Coastal (B), or Open Ocean (C). Pairwise comparisons of statistical significance between broad subclade classification means are represented in brackets above boxplots; 'ns' = no significance, '****' = p-value < 0.0001. D) World map showing the spatial distributions of the metagenomic samples used for recruitment to the SAR116 genomes. Every data point is a metagenomic sampling site, and marine environments are represented by color.

**Figure S6: Linear regression of log10-transformed subclade RPKM values vs. salinity values at all metagenomic sampling sites.** Data represented are from all three marine systems, and each panel represents a discrete SAR116 subclade. The trend line demonstrates the best fit from the quadratic regression model, and the shading around the trend line represents a 95% confidence interval of the fitted values. R-values quantify the correlation and directionality of the relationship between subclade RPKM and salinity.

**Figure S7: Linear regression of log10-transformed subclade RPKM values vs. salinity values at estuarine metagenomic sampling sites.** Data represented are from estuarine sites only, and each panel represents a discrete SAR116 subclade. The black trend line demonstrates the best fit from the quadratic regression model, and the shading around the trend line represents a 95% confidence interval of the fitted values. R-values quantify the correlation and directionality of the relationship between subclade RPKM and salinity.

**Figure S8: Linear regression of log10-transformed subclade RPKM values vs. salinity values at coastal metagenomic sampling sites.** Data represented are from coastal sites only, and each panel represents a discrete SAR116 subclade. The trend line demonstrates the best fit from the quadratic regression model, and the shading around the trend line represents a 95% confidence interval of the fitted values. R-values quantify the correlation and directionality of the relationship between subclade RPKM and salinity.

**Figure S9: Linear regression of log10-transformed subclade RPKM values vs. salinity values at open ocean metagenomic sampling sites.** Data represented are from open ocean sites only, and each panel represents a discrete SAR116 subclade. The trend line demonstrates the best fit from the quadratic regression model, and the shading around the trend line represents a 95% confidence interval of the fitted values. R-values quantify the correlation and directionality of the relationship between subclade RPKM and salinity.

**Figure S10: Linear regression of log10-transformed subclade RPKM values vs. temperature at all metagenomic sampling site.** Data represented are from all three marine systems, and each panel represents a discrete SAR116 subclade. The black trend line demonstrates the best fit from the quadratic regression model, and the shading around the trend line represents a 95% confidence interval of the fitted values. R-values quantify the correlation and directionality of the relationship between subclade RPKM and temperature.

**Figure S11: Linear regression of log10-transformed subclade RPKM values vs. temperature at coastal metagenomic sampling sites.** Data represented are from coastal sites only, and each panel represents a discrete SAR116 subclade. The black trend line demonstrates the best fit from the quadratic regression model, and the shading around the trend line represents a 95% confidence interval of the fitted values. R-values quantify the correlation and directionality of the relationship between subclade RPKM and temperature.

**Figure S12: Linear regression of log10-transformed subclade RPKM values vs. temperature at open ocean metagenomic sampling sites.** Data represented are from open ocean sites only, and each panel represents a discrete SAR116 subclade. The black trend line demonstrates the best fit from the quadratic regression model, and the shading around the trend line represents a 95% confidence interval of the fitted values. R-values quantify the correlation and directionality of the relationship between subclade RPKM and temperature.

**Figure S13: Quadratic regression of log10-transformed subclade RPKM values vs. temperature at estuarine metagenomic sampling sites.** Data represented are from estuarine sites only, and each panel represents a discrete SAR116 subclade. The black trend line demonstrates the best fit from the quadratic regression model, and the shading around the trend line represents a 95% confidence interval of the fitted values. R-values quantify the correlation and directionality of the relationship between subclade RPKM and temperature.

**Figure S14: Growth curves for LSUCC SAR116 isolates salinity tolerance.** Rows represent different salinity values in increasing order of salinity (ppt), and columns represent the different LSUCC isolates. Biological replicates are represented in each growth curve by different colors.

**Figure S15: Growth curves for LSUCC SAR116 isolates temperature tolerance.** Rows represent the different LSUCC isolates, and columns represent the different temperatures in increasing order of temperature (ºC). Biological replicates are represented in each growth curve by different colors.

**Figure S16: Panel of SEM image size measurements for LSUCC0719.** Measurements taken are indicated by the red lines on each of the panels, and the corresponding measurement is recorded in red text next to the line.

**Figure S17: Panel of SEM image size measurements for LSUCC0744.** Measurements taken are indicated by the red lines on each of the panels, and the corresponding measurement is recorded in red text next to the line.

**Figure S18: Panel of SEM image size measurements for LSUCC0684.** Measurements taken are indicated by the red lines on each of the panels, and the corresponding measurement is recorded in red text next to the line.

**Figure S19: Phylogenetic tree of SoxB**. S-sulfosulfanyl-L-cysteine sulfohydrolase, with closest relatives from NCBI RefSeq database, represented by black tips. Blue, green, orange, pink, and yellow tips indicate SAR116 SoxB protein sequences. Scale bar represents 0.08 changes per position. Bootstrap support values (n=1000) are indicated at nodes.

**Figure S20: Phylogenetic tree of FccB**. sulfide dehydrogenase [flavocytochrome c], with closest relatives from NCBI RefSeq database, represented by black tips. Blue, green, orange, pink, and yellow tips indicate SAR116 FccB protein sequences. Scale bar represents 1 change per position. Bootstrap support values (n=1000) are indicated at nodes.

**Figure S21: Phylogenetic tree of SoeA**. sulfite dehydrogenase (quinone), with closest relatives from NCBI RefSeq database, represented by black tips. Blue, green, purple and yellow tips indicate SAR116 SoeA protein sequences. Scale bar represents 0.1 changes per position. Bootstrap support values (n=1000) are indicated at nodes.

**Figure S21: Panel of LSUCC0684 (Calcibacteria) Pleomorphy**. Measurements taken are indicated by the red lines on each of the panels, and the corresponding measurement is recorded in red text next to the line. LSUCC0684 "calcibacteria" are represented by the small flattened circular cells.

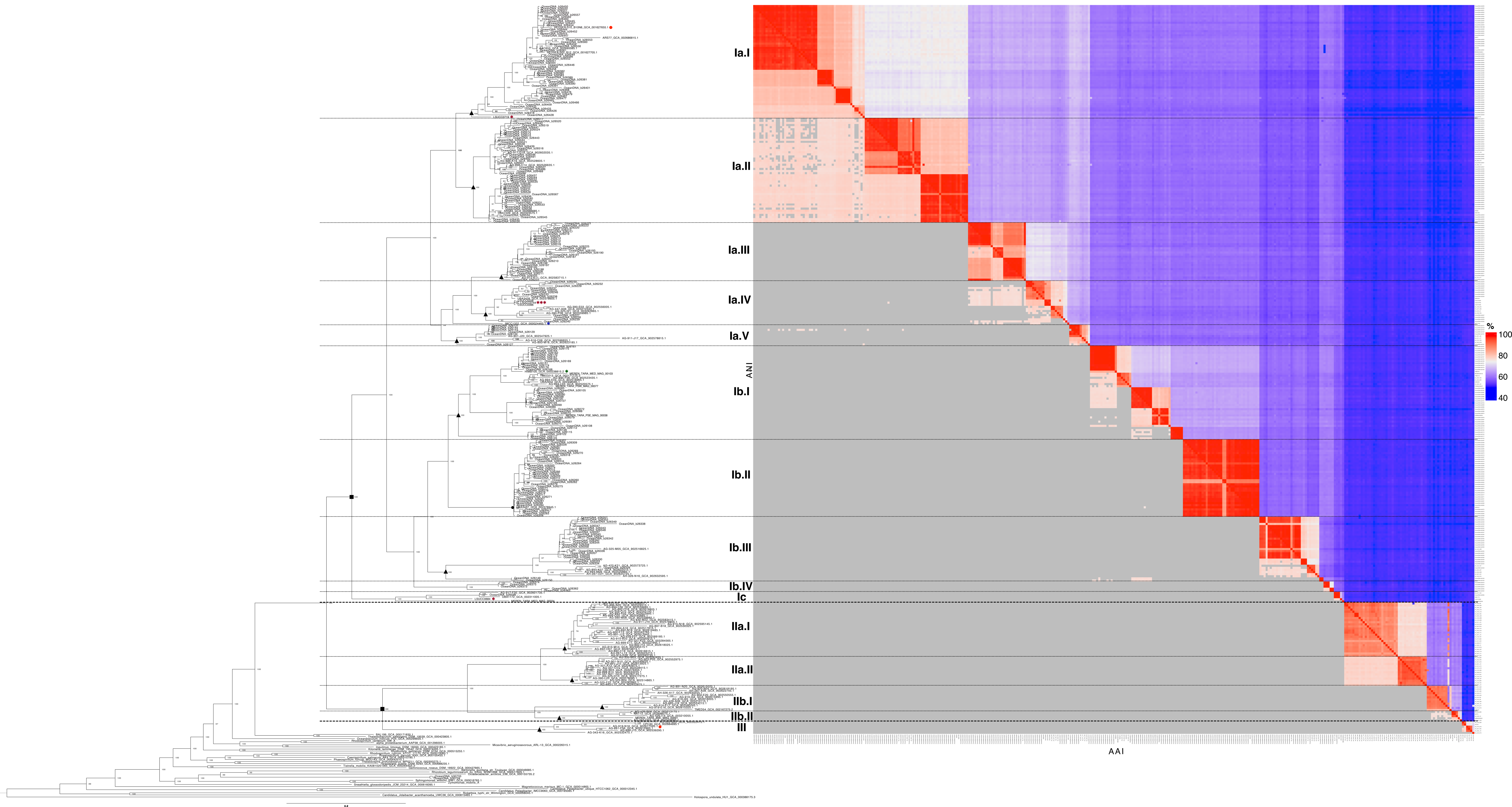

Ia.I
Ia.II
Ia.III
Ia.IV
Ia.V
Ib.I
Ib.II
Ib.III
Ib.IV
Ic
IIa.I
IIa.II
IIb.I
IIb.II
III
ANI
AAI
%
100
80
60
40

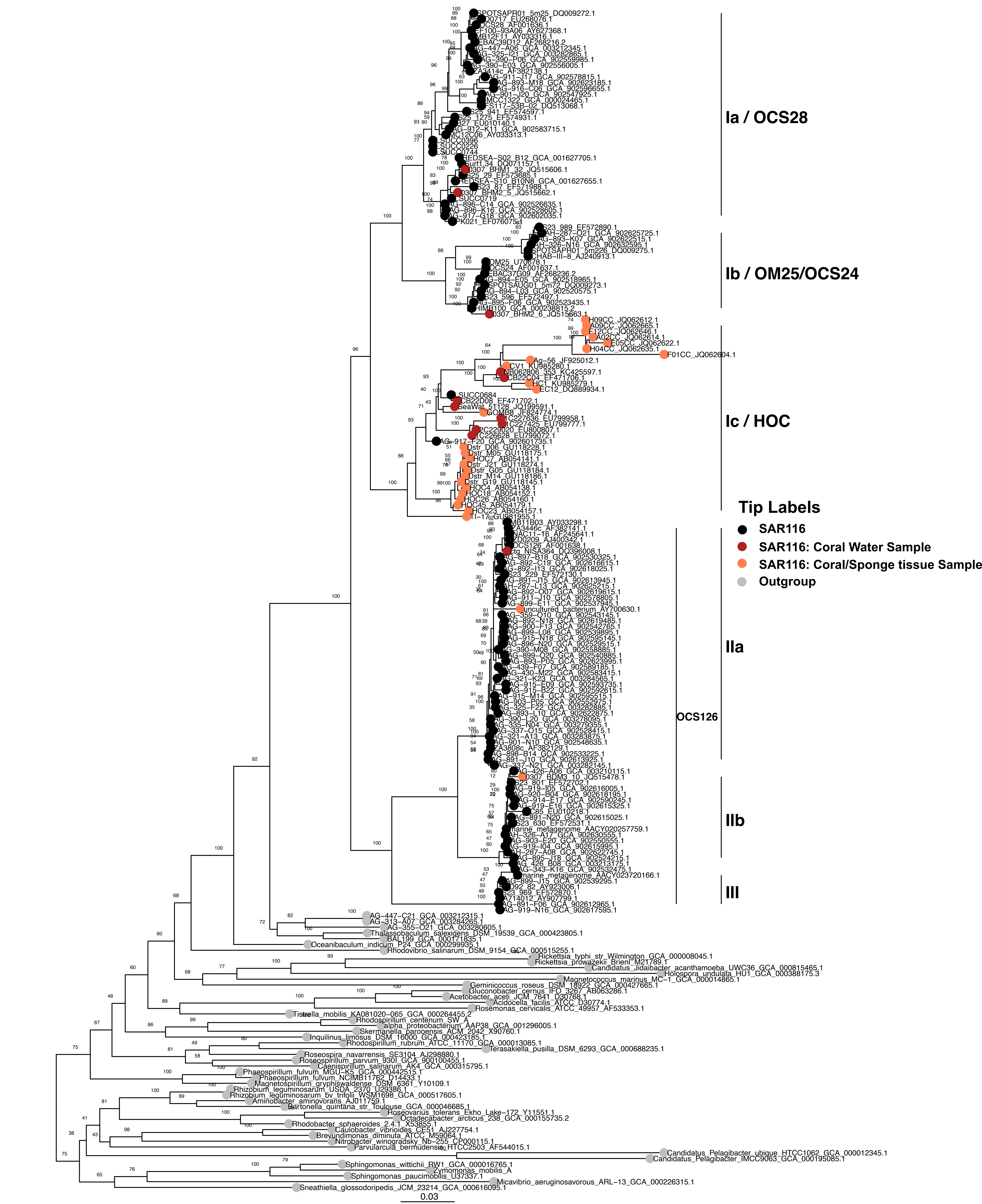
Ia / OCS28
Ib / OM25/OCS24
Ic / HOC
Tip Labels
SAR116
SAR116: Coral Water Sample
SAR116: Coral/Sponge tissue Sample
Outgroup
IIa
OCS126
IIb
III
0.03

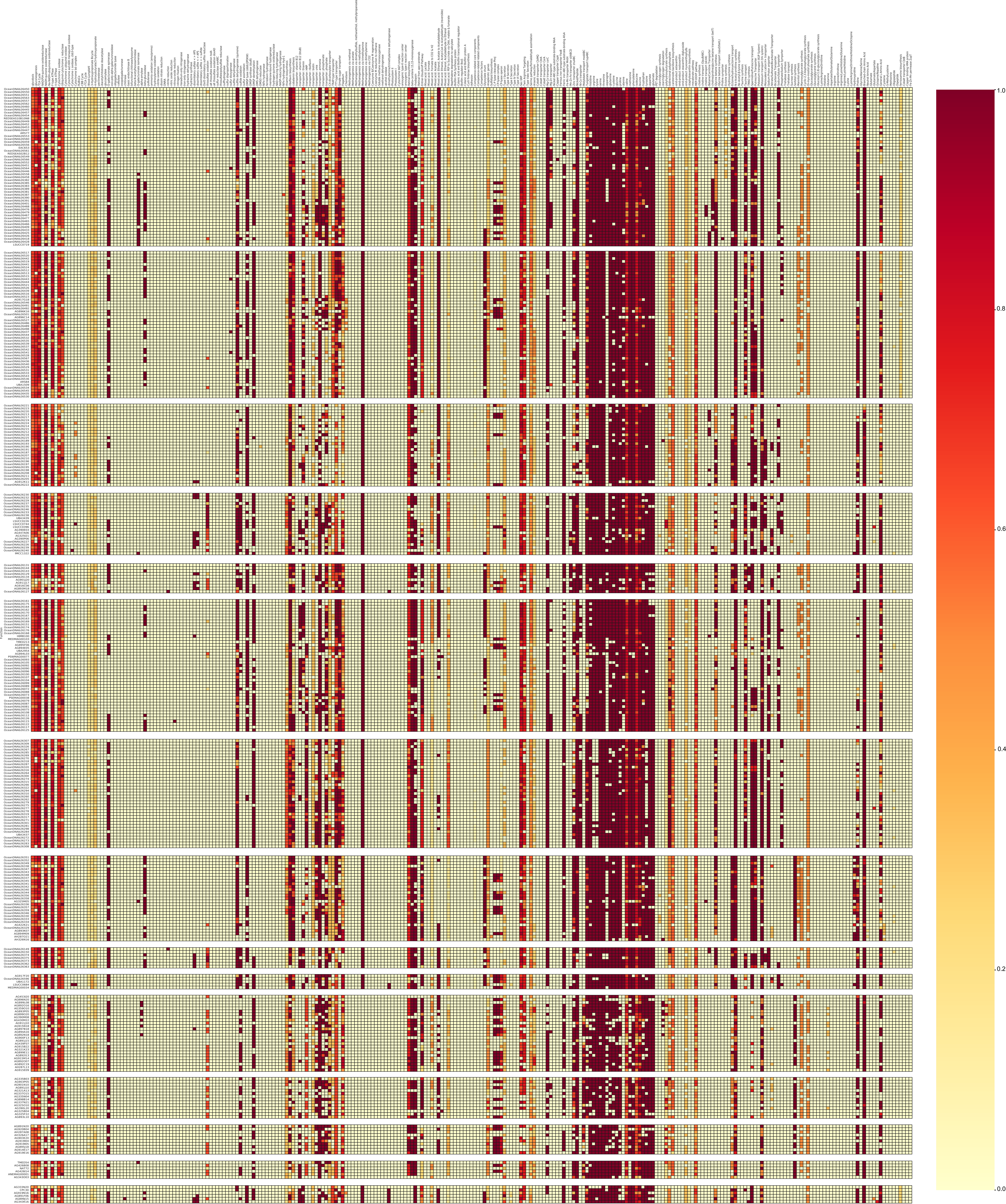
1.0
0.8
0.6
0.4
0.2
0.0

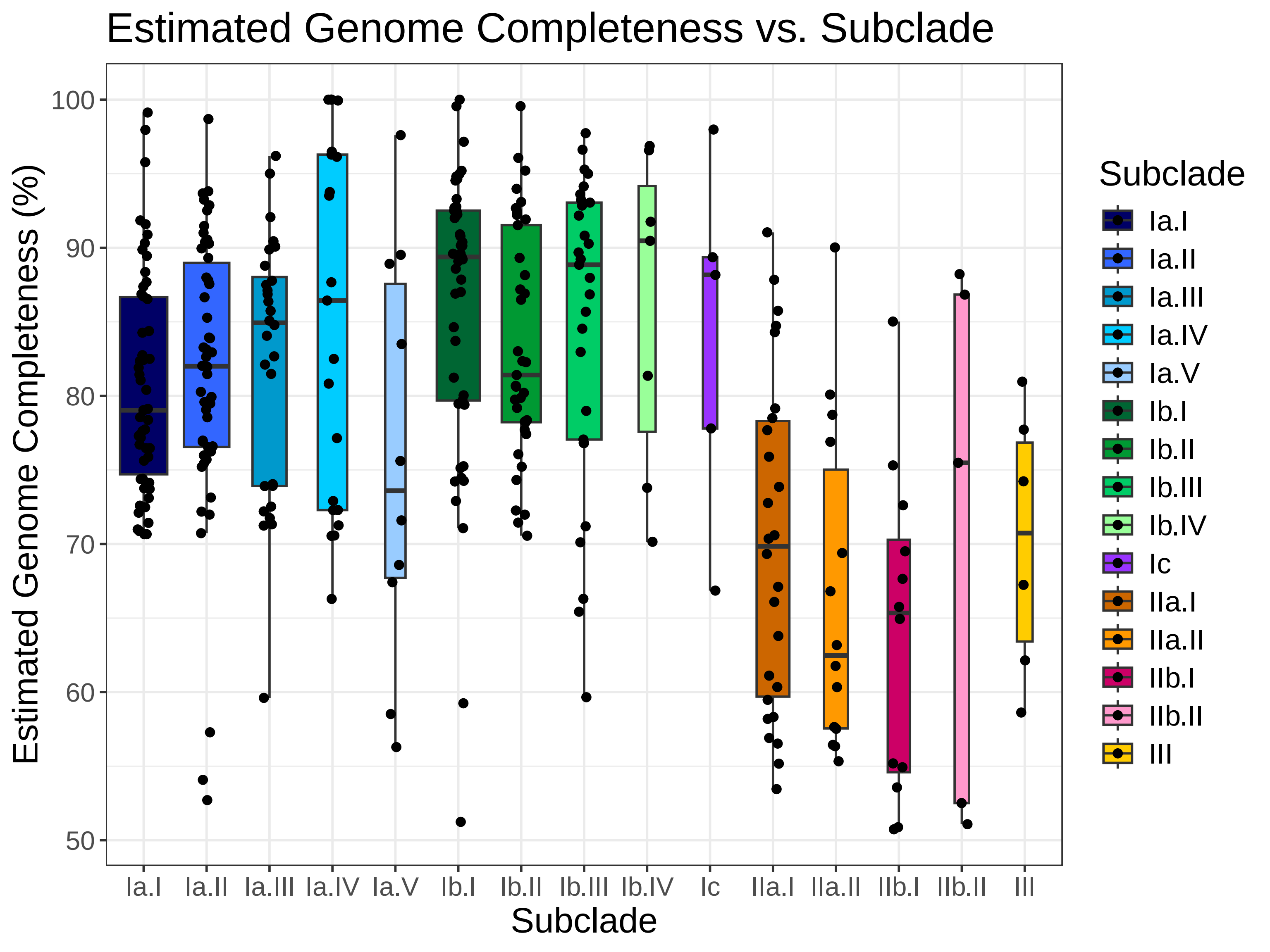

Estimated Genome Completeness vs. Subclade
Estimated Genome Completeness (%)
100
90
80
70
60
50
Ia.I
Ia.II
Ia.III
Ia.IV
Ia.V
Ib.I
Ib.II
Ib.III
Ib.IV
Ic
IIa.I
IIa.II
IIb.I
IIb.II
III
Subclade
Subclade
Ia.I
Ia.II
Ia.III
Ia.IV
Ia.V
Ib.I
Ib.II
Ib.III
Ib.IV
Ic
IIa.I
IIa.II
IIb.I
IIb.II
III

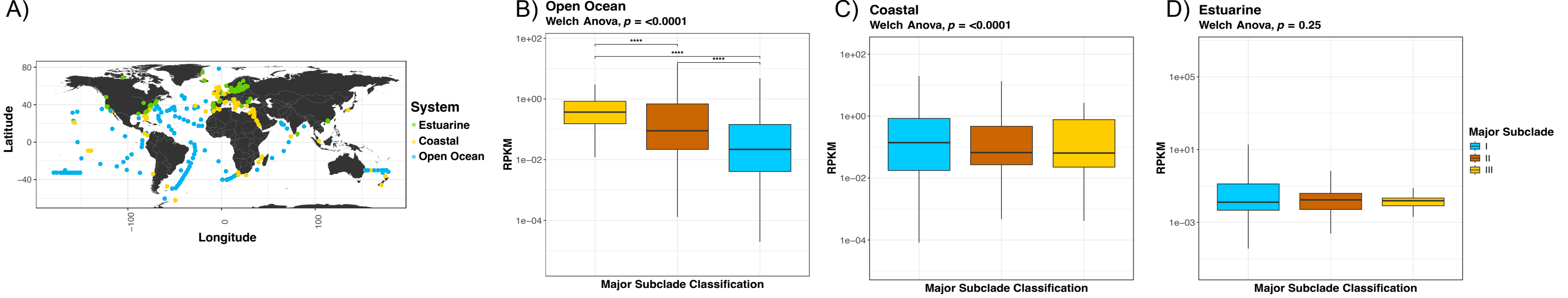

A)
Latitude
80
40
0
−40
−100
0
100
Longitude
System
Estuarine
Coastal
Open Ocean
B) Open Ocean
Welch Anova, p = <0.0001
****
****
****
1e+02
1e+00
1e−02
1e−04
RPKM
Major Subclade Classification
C) Coastal
Welch Anova, p = <0.0001
1e+02
1e+00
1e−02
1e−04
RPKM
Major Subclade Classification
D) Estuarine
Welch Anova, p = 0.25
1e+05
1e+01
1e−03
RPKM
Major Subclade Classification
Major Subclade
I
II
III

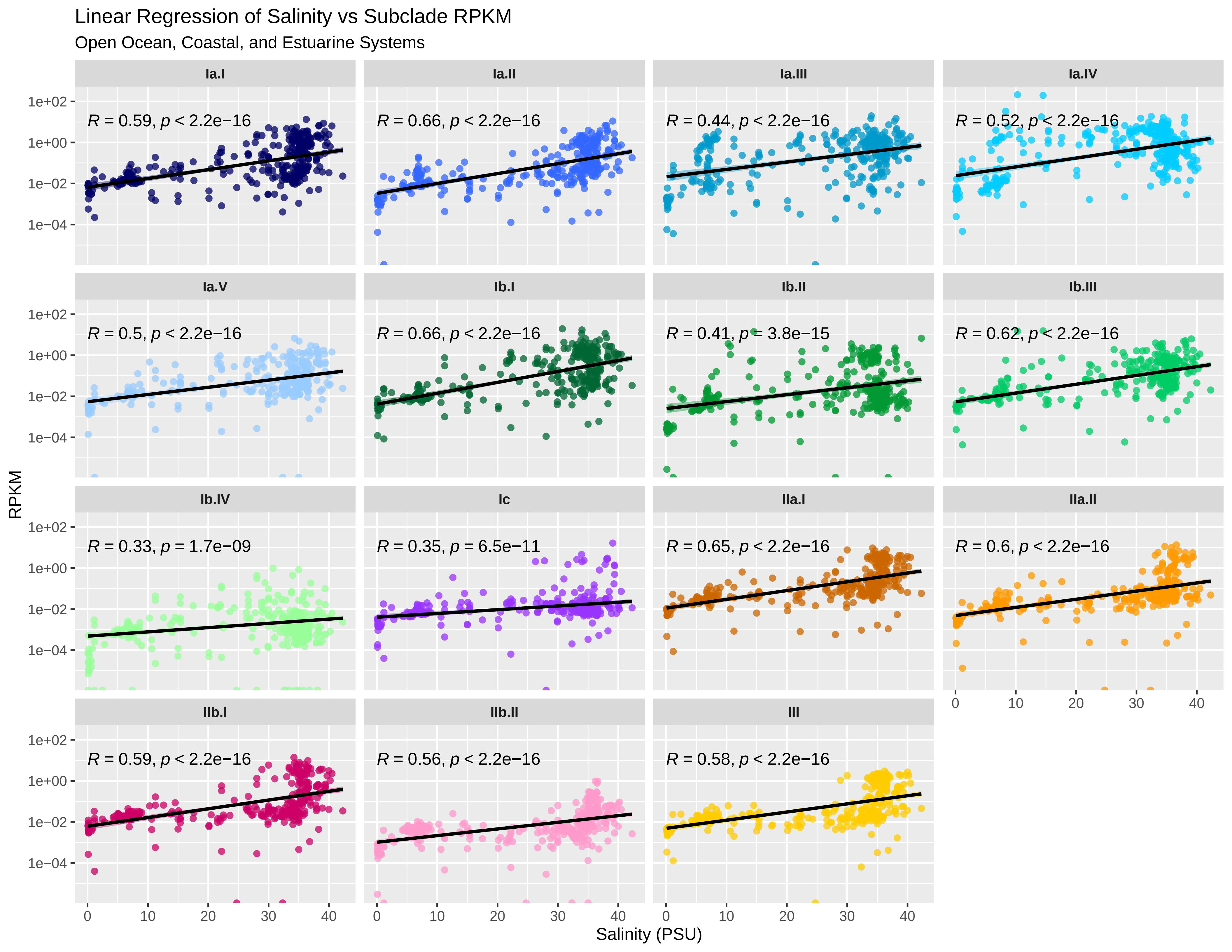
Linear Regression of Salinity vs Subclade RPKM
Open Ocean, Coastal, and Estuarine Systems
Ia.I
R = 0.59, p < 2.2e−16
Ia.II
R = 0.66, p < 2.2e−16
Ia.III
R = 0.44, p < 2.2e−16
Ia.IV
R = 0.52, p < 2.2e−16
Ia.V
R = 0.5, p < 2.2e−16
Ib.I
R = 0.66, p < 2.2e−16
Ib.II
R = 0.41, p = 3.8e−15
Ib.III
R = 0.62, p < 2.2e−16
Ib.IV
R = 0.33, p = 1.7e−09
Ic
R = 0.35, p = 6.5e−11
IIa.I
R = 0.65, p < 2.2e−16
IIa.II
R = 0.6, p < 2.2e−16
IIb.I
R = 0.59, p < 2.2e−16
IIb.II
R = 0.56, p < 2.2e−16
III
R = 0.58, p < 2.2e−16
1e+02
1e+00
1e−02
1e−04
0
10
20
30
40
RPKM
Salinity (PSU)

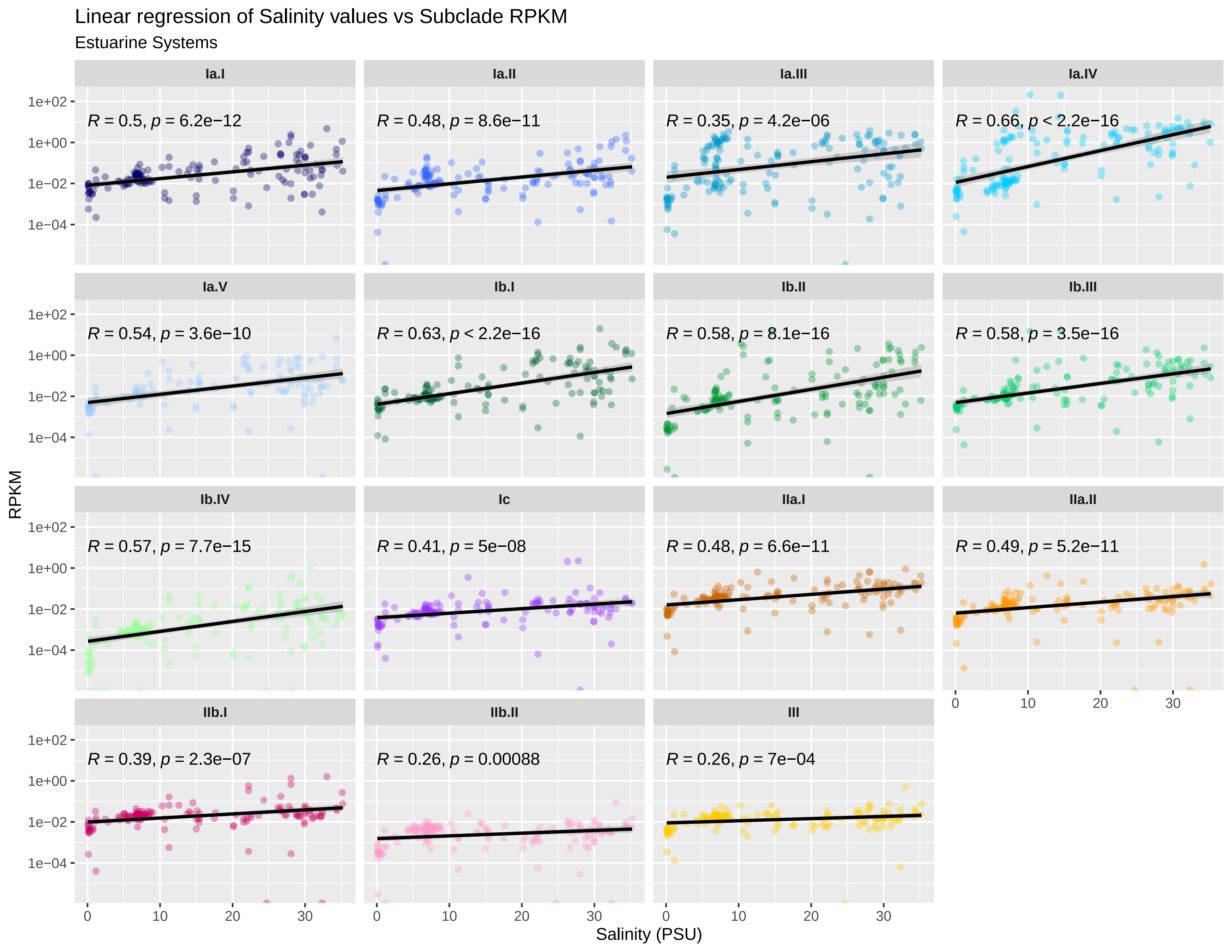

Linear regression of Salinity values vs Subclade RPKM
Estuarine Systems
Ia.I
R = 0.5, p = 6.2e−12
Ia.II
R = 0.48, p = 8.6e−11
Ia.III
R = 0.35, p = 4.2e−06
Ia.IV
R = 0.66, p < 2.2e−16
Ia.V
R = 0.54, p = 3.6e−10
Ib.I
R = 0.63, p < 2.2e−16
Ib.II
R = 0.58, p = 8.1e−16
Ib.III
R = 0.58, p = 3.5e−16
Ib.IV
R = 0.57, p = 7.7e−15
Ic
R = 0.41, p = 5e−08
IIa.I
R = 0.48, p = 6.6e−11
IIa.II
R = 0.49, p = 5.2e−11
IIb.I
R = 0.39, p = 2.3e−07
IIb.II
R = 0.26, p = 0.00088
III
R = 0.26, p = 7e−04
RPKM
1e+02
1e+00
1e−02
1e−04
0
10
20
30
Salinity (PSU)

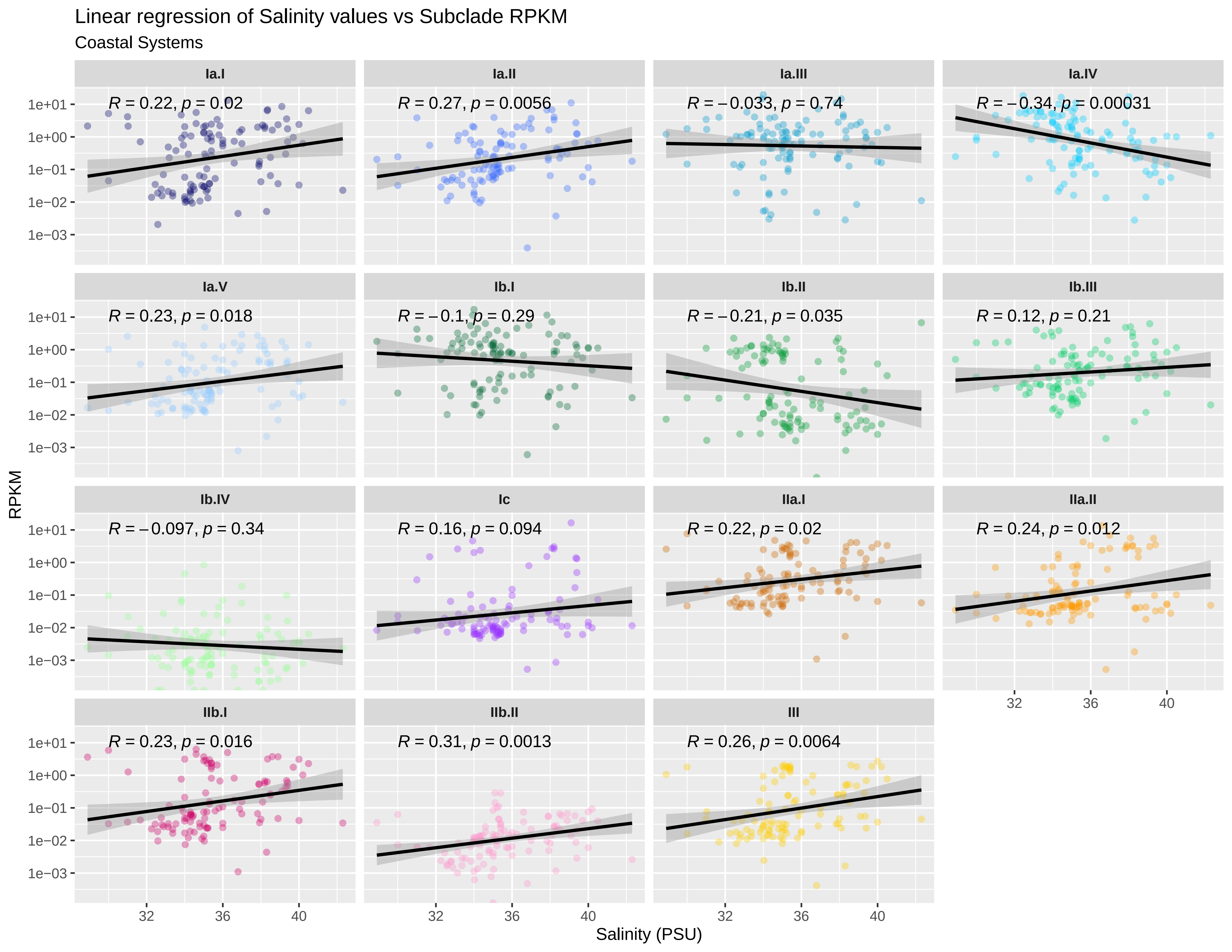

Linear regression of Salinity values vs Subclade RPKM
Coastal Systems
Ia.I
R = 0.22, p = 0.02
Ia.II
R = 0.27, p = 0.0056
Ia.III
R = −0.033, p = 0.74
Ia.IV
R = −0.34, p = 0.00031
Ia.V
R = 0.23, p = 0.018
Ib.I
R = −0.1, p = 0.29
Ib.II
R = −0.21, p = 0.035
Ib.III
R = 0.12, p = 0.21
Ib.IV
R = −0.097, p = 0.34
Ic
R = 0.16, p = 0.094
IIa.I
R = 0.22, p = 0.02
IIa.II
R = 0.24, p = 0.012
IIb.I
R = 0.23, p = 0.016
IIb.II
R = 0.31, p = 0.0013
III
R = 0.26, p = 0.0064
1e+01
1e+00
1e−01
1e−02
1e−03
32
36
40
RPKM
Salinity (PSU)

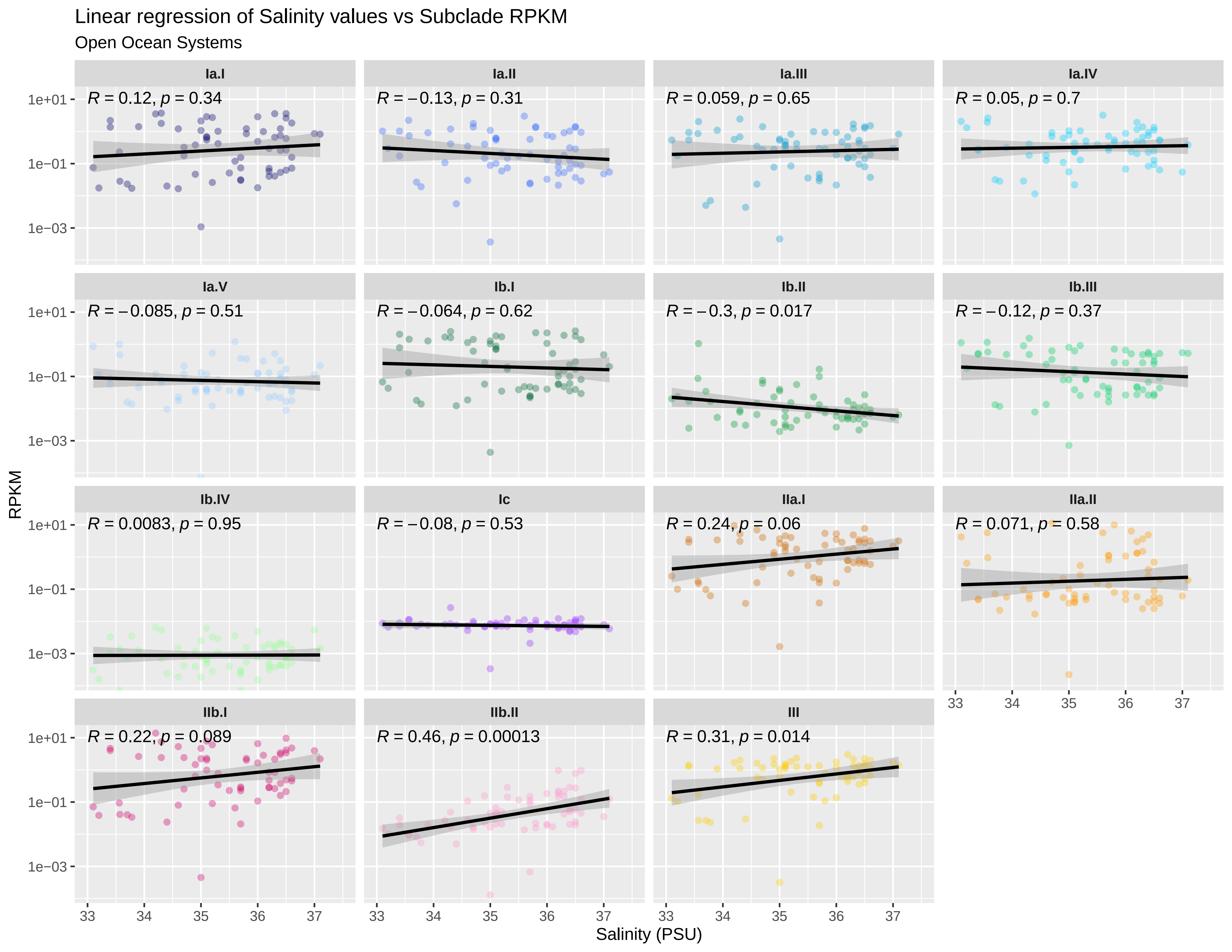
Linear regression of Salinity values vs Subclade RPKM
Open Ocean Systems
Ia.I
R = 0.12, p = 0.34
Ia.II
R = –0.13, p = 0.31
Ia.III
R = 0.059, p = 0.65
Ia.IV
R = 0.05, p = 0.7
Ia.V
R = –0.085, p = 0.51
Ib.I
R = –0.064, p = 0.62
Ib.II
R = –0.3, p = 0.017
Ib.III
R = –0.12, p = 0.37
Ib.IV
R = 0.0083, p = 0.95
Ic
R = –0.08, p = 0.53
IIa.I
R = 0.24, p = 0.06
IIa.II
R = 0.071, p = 0.58
IIb.I
R = 0.22, p = 0.089
IIb.II
R = 0.46, p = 0.00013
III
R = 0.31, p = 0.014
1e+01
1e–01
1e–03
RPKM
33
34
35
36
37
Salinity (PSU)

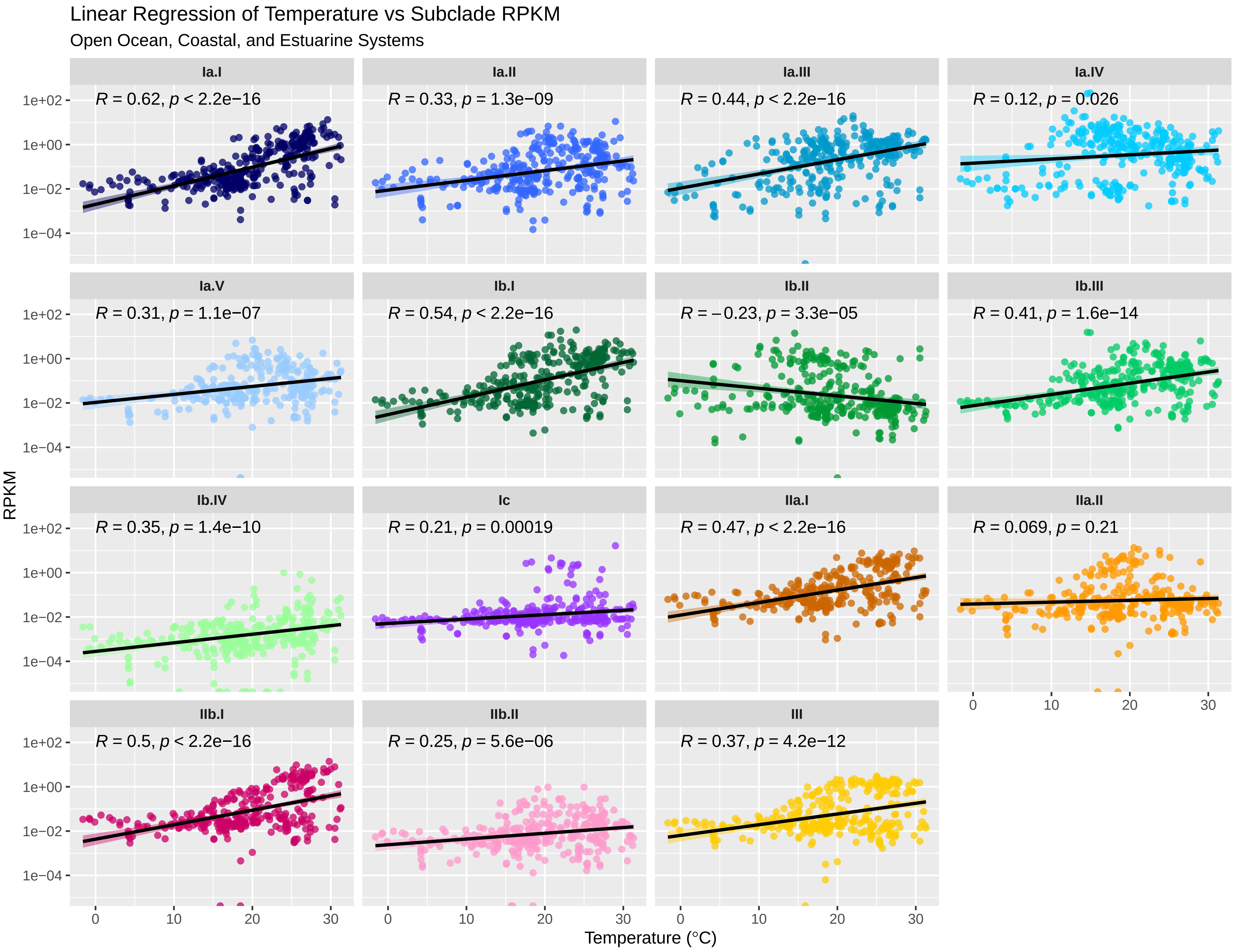

Linear Regression of Temperature vs Subclade RPKM
Open Ocean, Coastal, and Estuarine Systems
Ia.I
R = 0.62, p < 2.2e−16
Ia.II
R = 0.33, p = 1.3e−09
Ia.III
R = 0.44, p < 2.2e−16
Ia.IV
R = 0.12, p = 0.026
Ia.V
R = 0.31, p = 1.1e−07
Ib.I
R = 0.54, p < 2.2e−16
Ib.II
R = −0.23, p = 3.3e−05
Ib.III
R = 0.41, p = 1.6e−14
Ib.IV
R = 0.35, p = 1.4e−10
Ic
R = 0.21, p = 0.00019
IIa.I
R = 0.47, p < 2.2e−16
IIa.II
R = 0.069, p = 0.21
IIb.I
R = 0.5, p < 2.2e−16
IIb.II
R = 0.25, p = 5.6e−06
III
R = 0.37, p = 4.2e−12
RPKM
1e+02
1e+00
1e−02
1e−04
0
10
20
30
Temperature (°C)

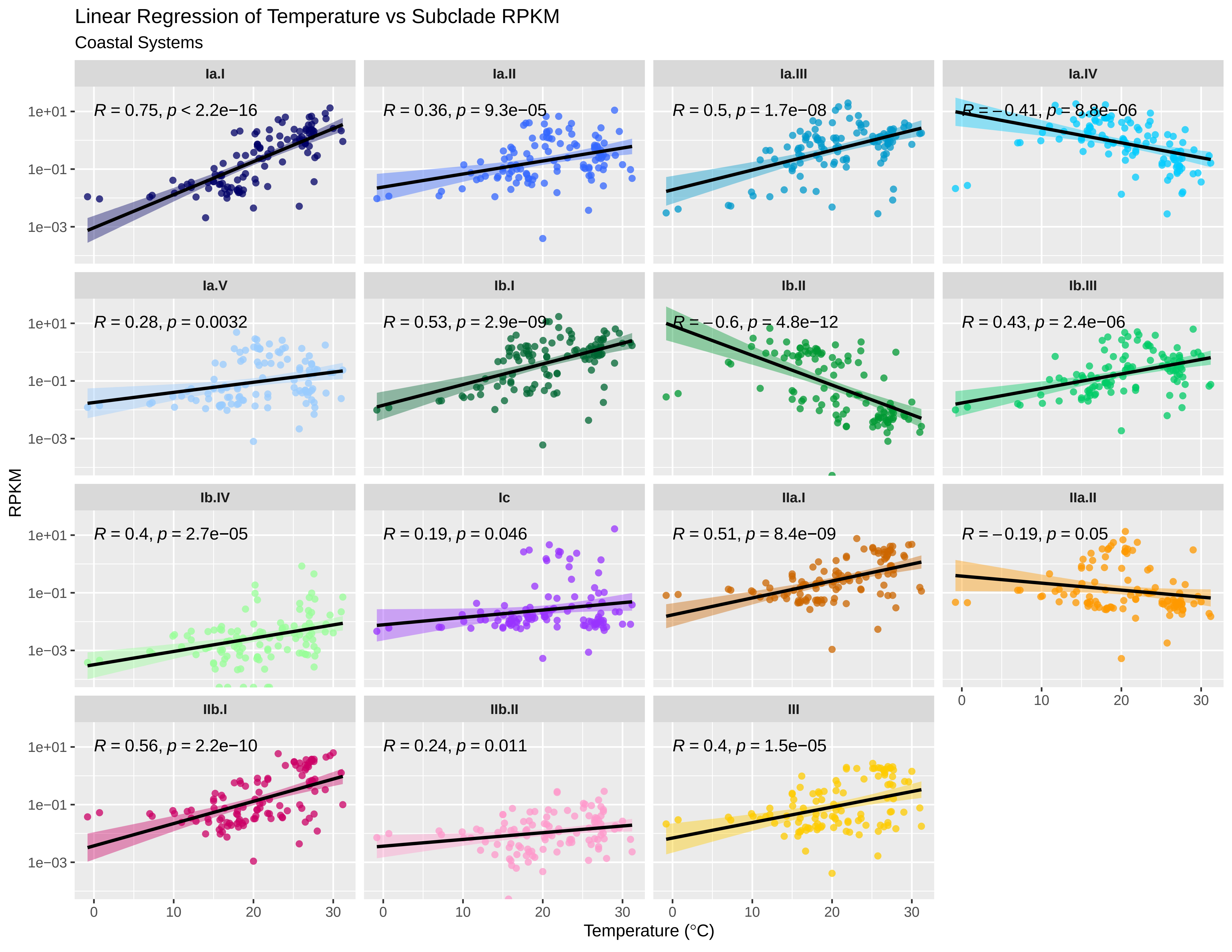
Linear Regression of Temperature vs Subclade RPKM
Coastal Systems
Ia.I
R = 0.75, p < 2.2e−16
Ia.II
R = 0.36, p = 9.3e−05
Ia.III
R = 0.5, p = 1.7e−08
Ia.IV
R = −0.41, p = 8.8e−06
Ia.V
R = 0.28, p = 0.0032
Ib.I
R = 0.53, p = 2.9e−09
Ib.II
R = −0.6, p = 4.8e−12
Ib.III
R = 0.43, p = 2.4e−06
Ib.IV
R = 0.4, p = 2.7e−05
Ic
R = 0.19, p = 0.046
IIa.I
R = 0.51, p = 8.4e−09
IIa.II
R = −0.19, p = 0.05
IIb.I
R = 0.56, p = 2.2e−10
IIb.II
R = 0.24, p = 0.011
III
R = 0.4, p = 1.5e−05
RPKM
1e+01
1e−01
1e−03
0
10
20
30
Temperature (°C)

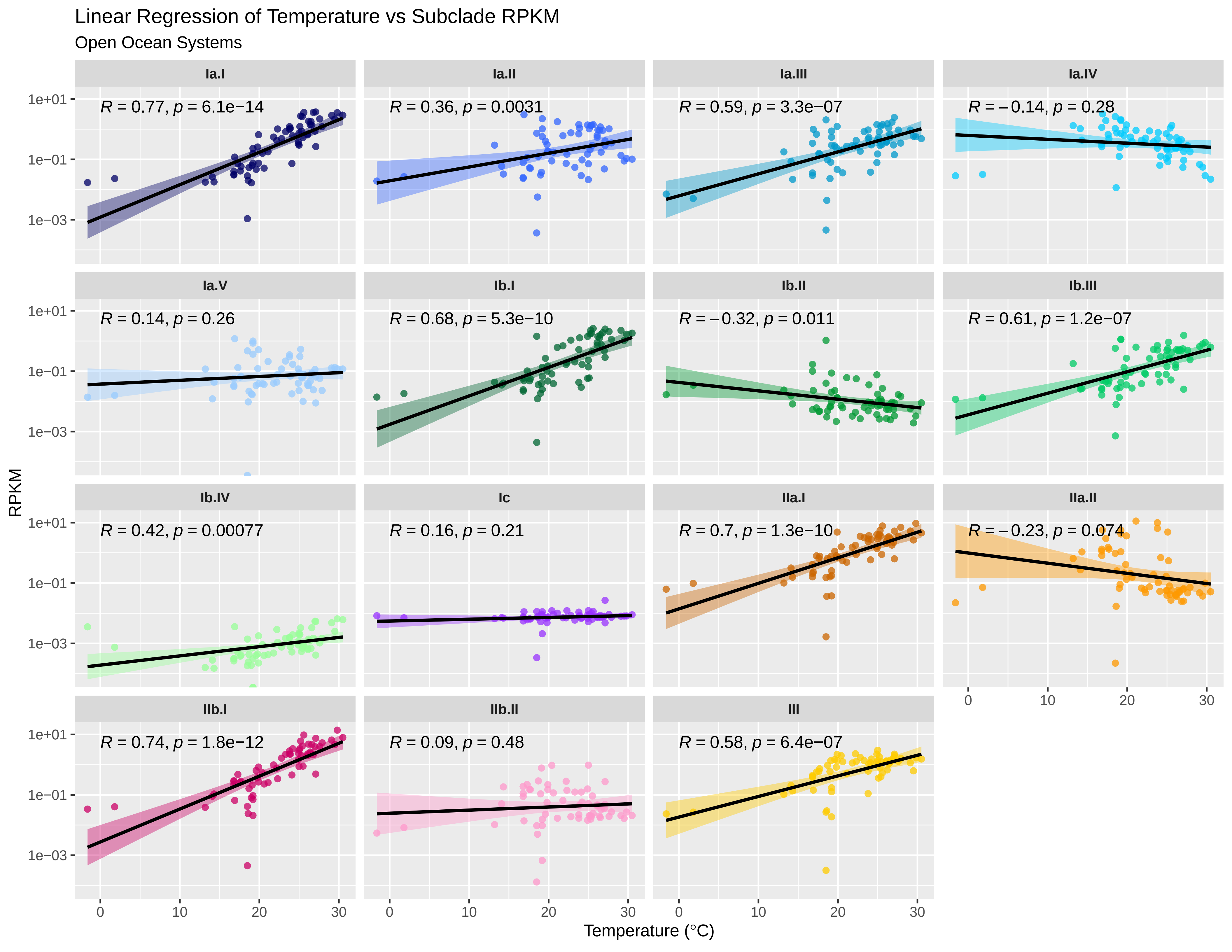

Linear Regression of Temperature vs Subclade RPKM
Open Ocean Systems
Ia.I
R = 0.77, p = 6.1e−14
Ia.II
R = 0.36, p = 0.0031
Ia.III
R = 0.59, p = 3.3e−07
Ia.IV
R = −0.14, p = 0.28
Ia.V
R = 0.14, p = 0.26
Ib.I
R = 0.68, p = 5.3e−10
Ib.II
R = −0.32, p = 0.011
Ib.III
R = 0.61, p = 1.2e−07
Ib.IV
R = 0.42, p = 0.00077
Ic
R = 0.16, p = 0.21
IIa.I
R = 0.7, p = 1.3e−10
IIa.II
R = −0.23, p = 0.074
IIb.I
R = 0.74, p = 1.8e−12
IIb.II
R = 0.09, p = 0.48
III
R = 0.58, p = 6.4e−07
RPKM
1e+01
1e−01
1e−03
0
10
20
30
Temperature (°C)

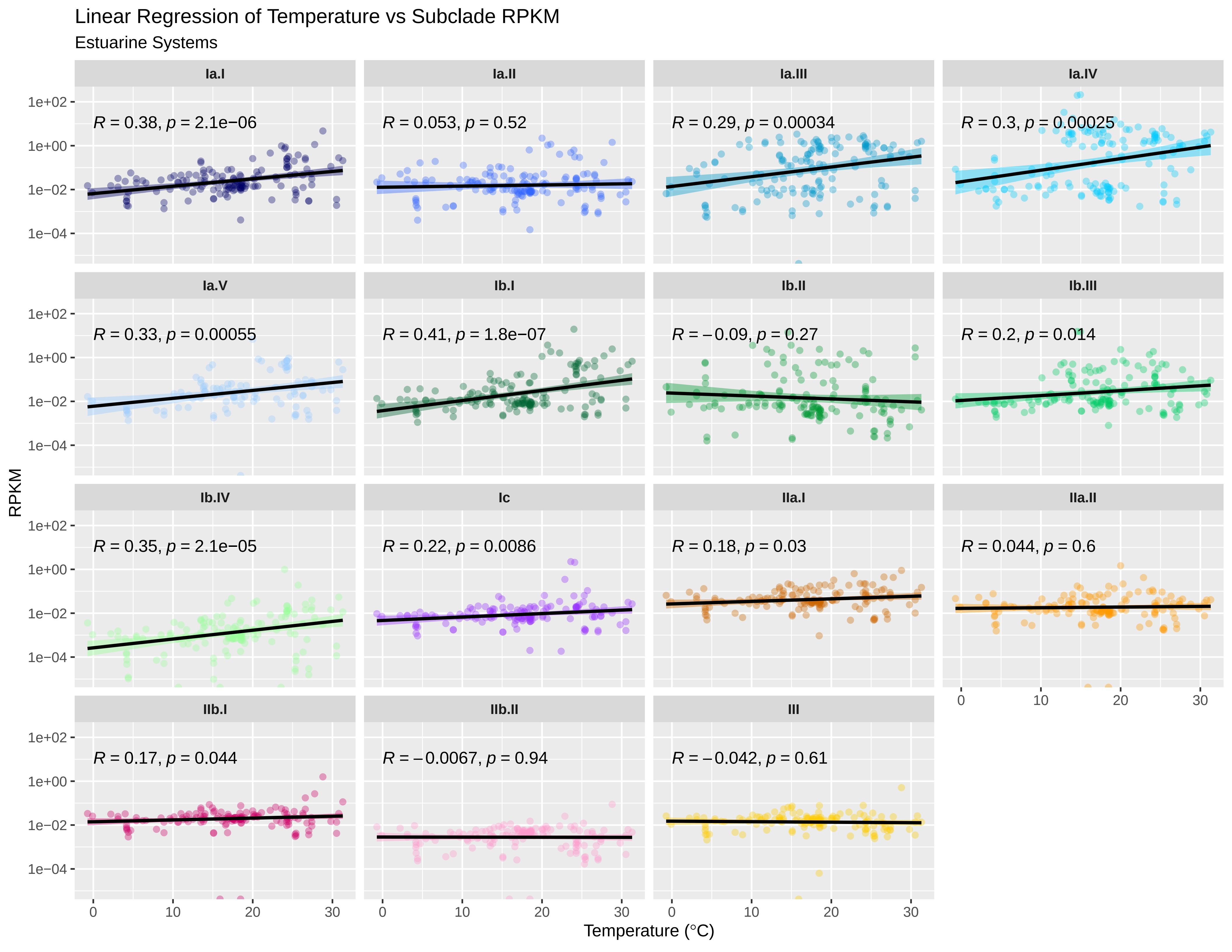

Linear Regression of Temperature vs Subclade RPKM
Estuarine Systems
Ia.I
R = 0.38, p = 2.1e−06
Ia.II
R = 0.053, p = 0.52
Ia.III
R = 0.29, p = 0.00034
Ia.IV
R = 0.3, p = 0.00025
Ia.V
R = 0.33, p = 0.00055
Ib.I
R = 0.41, p = 1.8e−07
Ib.II
R = −0.09, p = 0.27
Ib.III
R = 0.2, p = 0.014
Ib.IV
R = 0.35, p = 2.1e−05
Ic
R = 0.22, p = 0.0086
IIa.I
R = 0.18, p = 0.03
IIa.II
R = 0.044, p = 0.6
IIb.I
R = 0.17, p = 0.044
IIb.II
R = −0.0067, p = 0.94
III
R = −0.042, p = 0.61
RPKM
1e+02
1e+00
1e−02
1e−04
0
10
20
30
Temperature (°C)

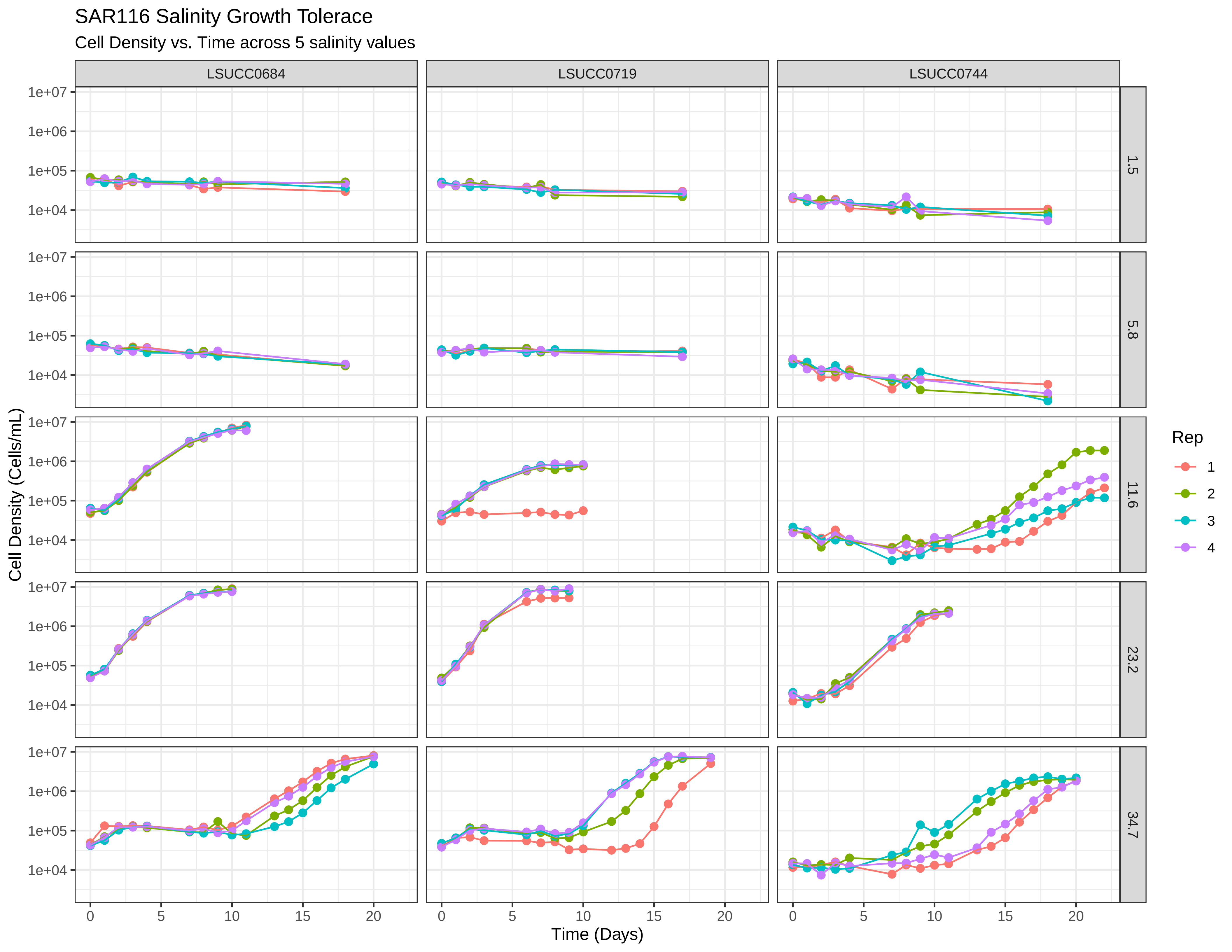
SAR116 Salinity Growth Tolerace
Cell Density vs. Time across 5 salinity values
LSUCC0684
LSUCC0719
LSUCC0744
1.5
5.8
11.6
23.2
34.7
1e+07
1e+06
1e+05
1e+04
Cell Density (Cells/mL)
0
5
10
15
20
Time (Days)
Rep
1
2
3
4

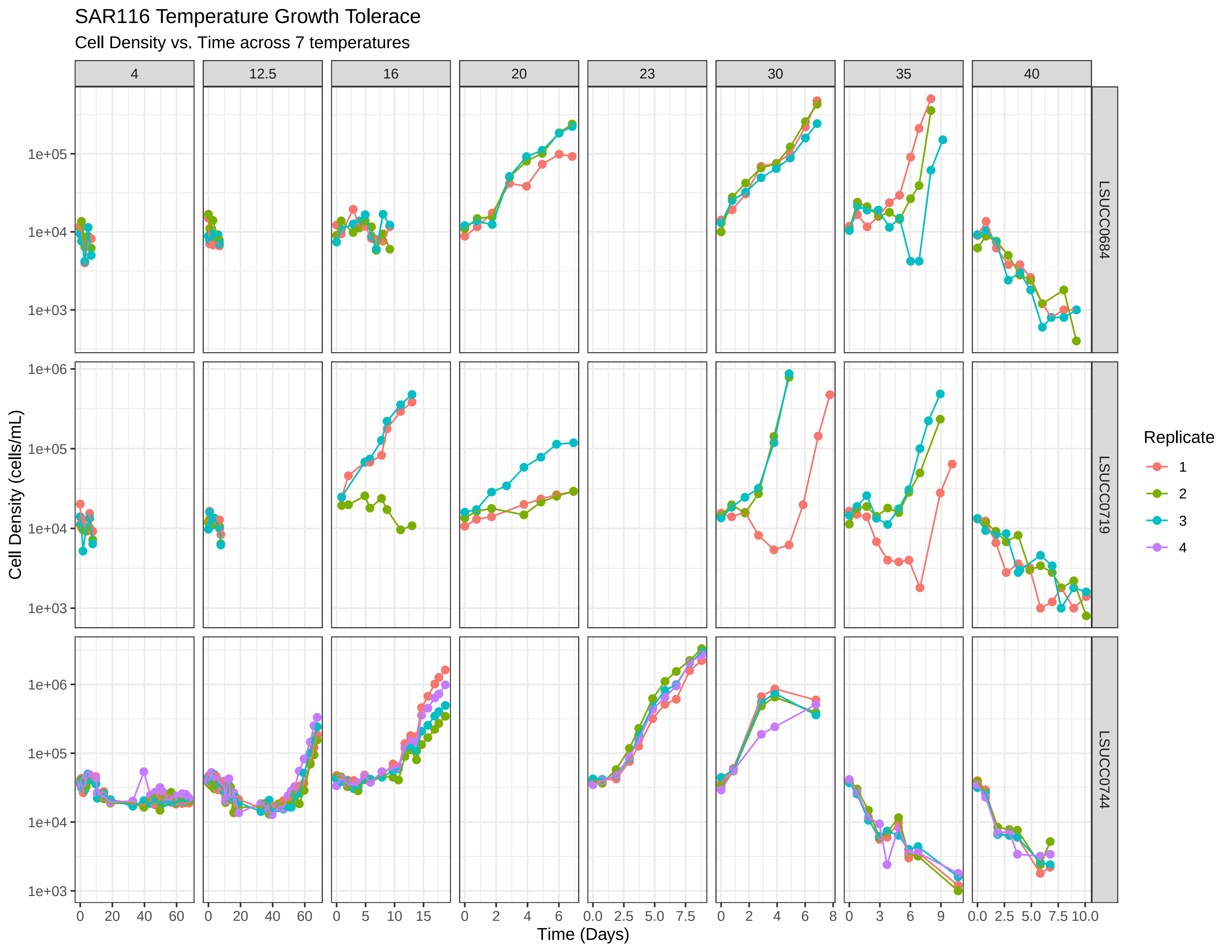
SAR116 Temperature Growth Tolerace
Cell Density vs. Time across 7 temperatures
4
12.5
16
20
23
30
35
40
LSUCC0684
LSUCC0719
LSUCC0744
Cell Density (cells/mL)
1e+03
1e+04
1e+05
1e+06
Time (Days)
Replicate
1
2
3
4

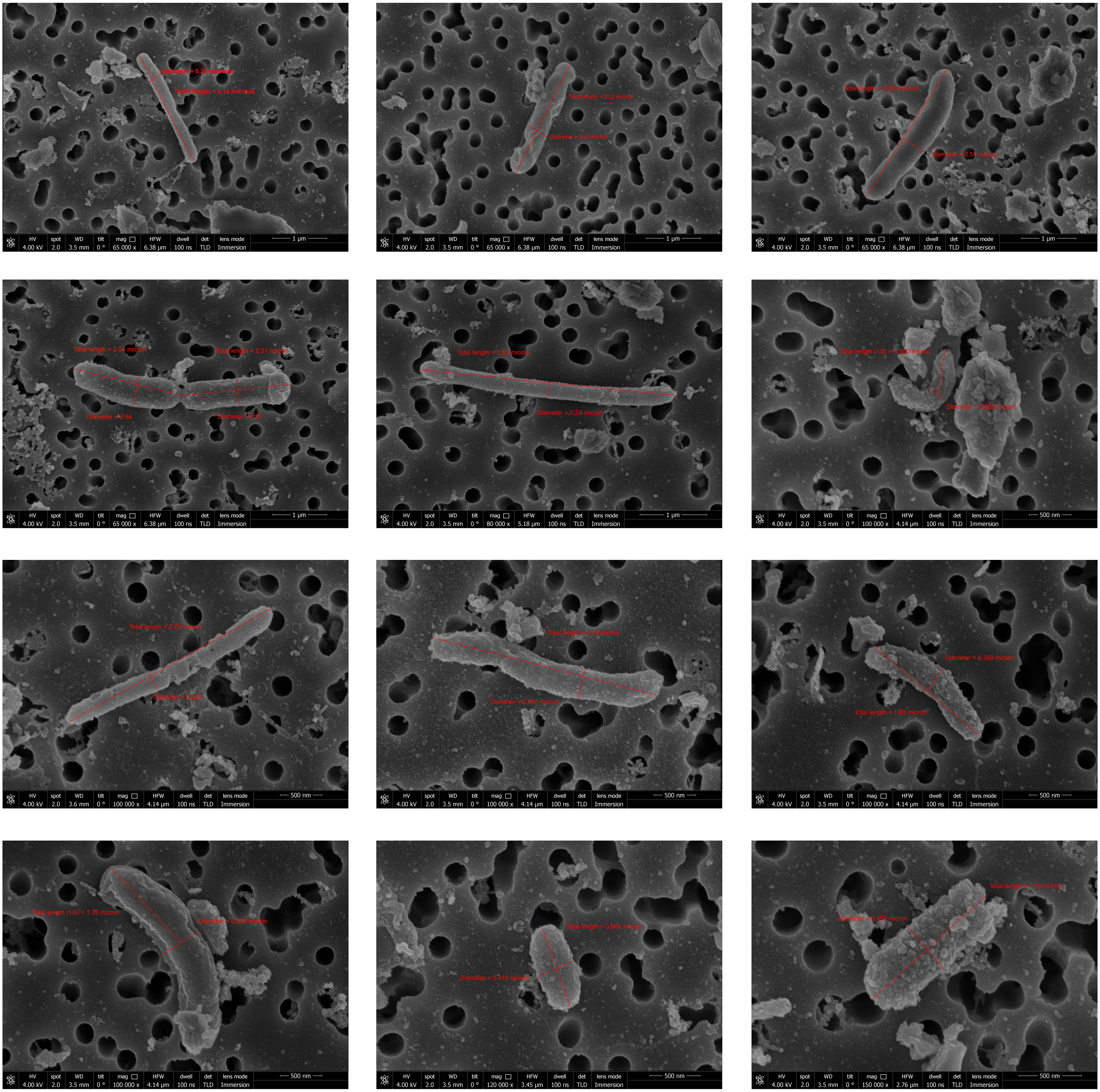

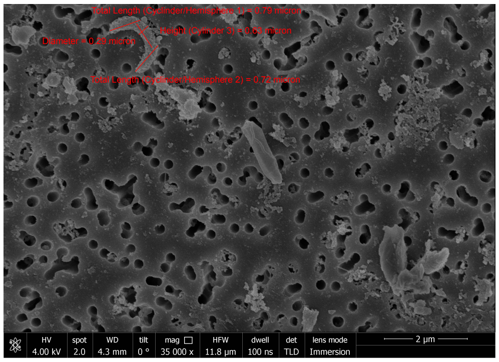
HV spot WD tilt mag HFW dwell det lens mode
4.00 kV 2.0 4.3 mm 0 ° 35 000 x 11.8 µm 100 ns TLD Immersion
2 µm
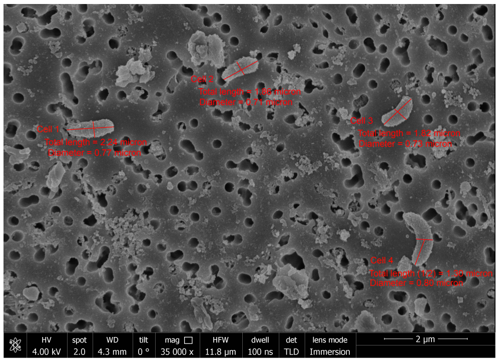
HV spot WD tilt mag HFW dwell det lens mode
4.00 kV 2.0 4.3 mm 0 ° 35 000 x 11.8 µm 100 ns TLD Immersion
2 µm
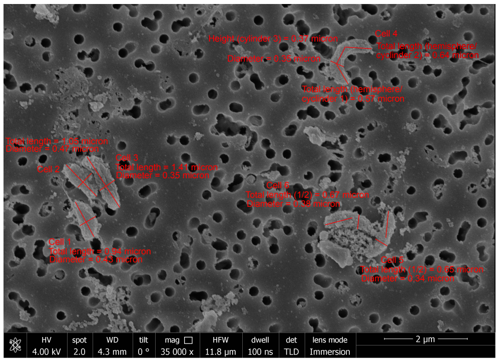
HV spot WD tilt mag HFW dwell det lens mode
4.00 kV 2.0 4.3 mm 0 ° 35 000 x 11.8 µm 100 ns TLD Immersion
2 µm
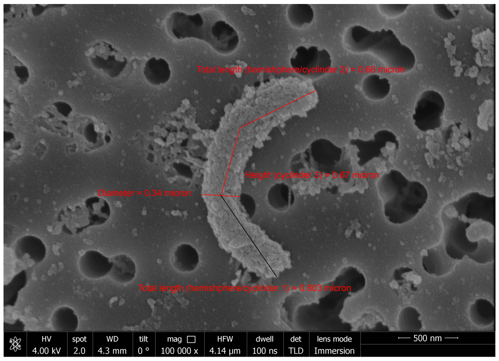
HV spot WD tilt mag HFW dwell det lens mode
4.00 kV 2.0 4.3 mm 0 ° 100 000 x 4.14 µm 100 ns TLD Immersion
500 nm
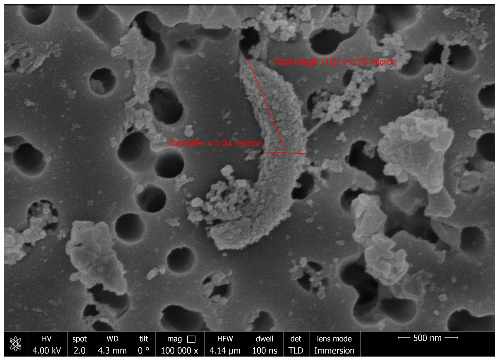
HV spot WD tilt mag HFW dwell det lens mode
4.00 kV 2.0 4.3 mm 0 ° 100 000 x 4.14 µm 100 ns TLD Immersion
500 nm
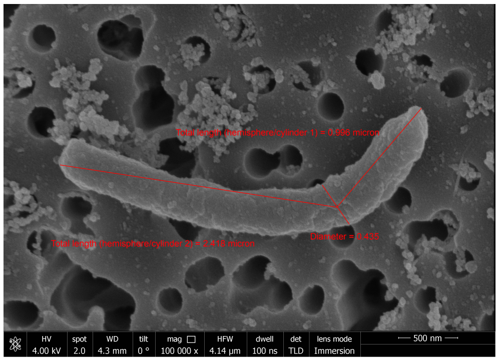
HV spot WD tilt mag HFW dwell det lens mode
4.00 kV 2.0 4.3 mm 0 ° 100 000 x 4.14 µm 100 ns TLD Immersion
500 nm
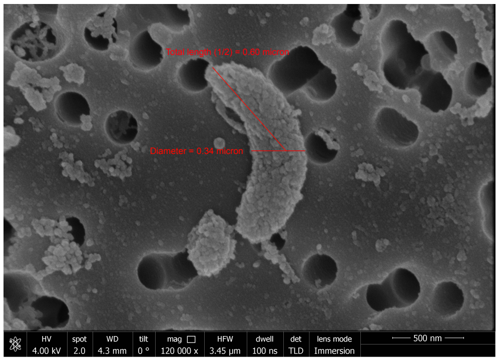
HV spot WD tilt mag HFW dwell det lens mode
4.00 kV 2.0 4.3 mm 0 ° 120 000 x 3.45 µm 100 ns TLD Immersion
500 nm
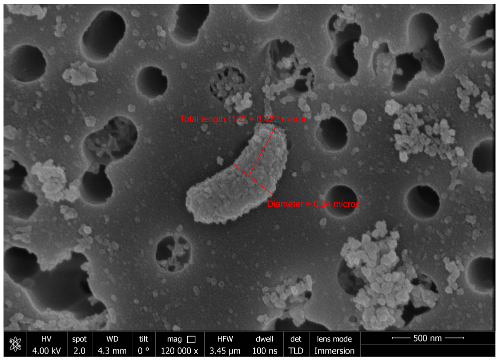
HV spot WD tilt mag HFW dwell det lens mode
4.00 kV 2.0 4.3 mm 0 ° 120 000 x 3.45 µm 100 ns TLD Immersion
500 nm
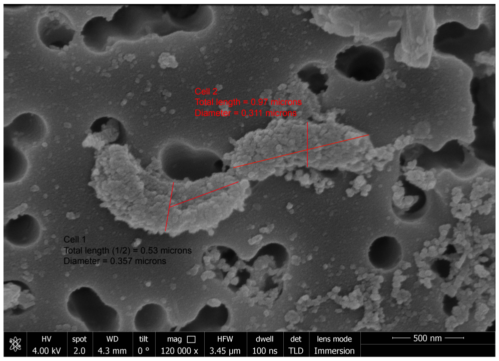
Cell 1
Total length (1/2) = 0.53 microns
Diameter = 0.357 microns
HV spot WD tilt mag HFW dwell det lens mode
4.00 kV 2.0 4.3 mm 0 ° 120 000 x 3.45 µm 100 ns TLD Immersion
500 nm

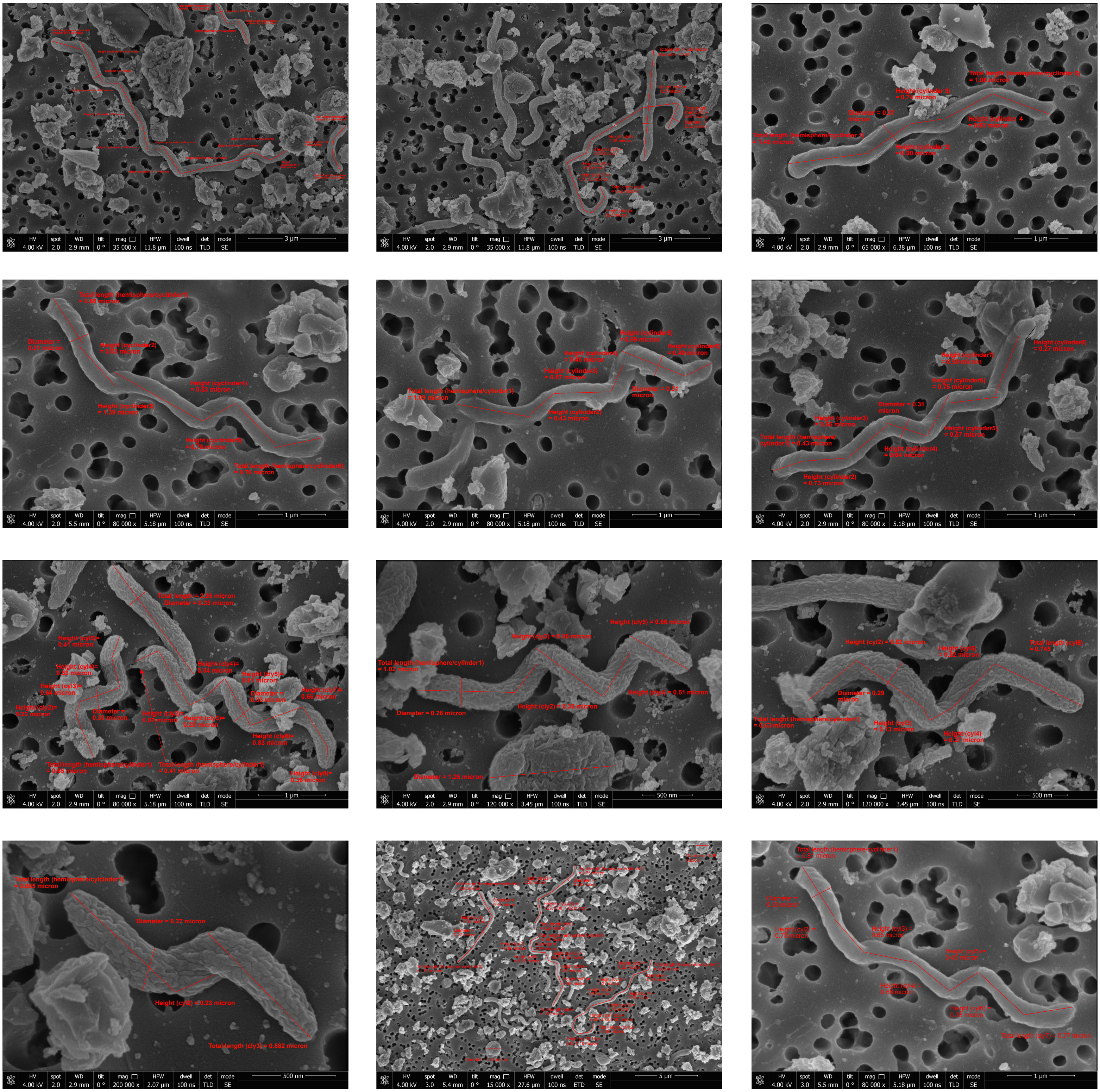

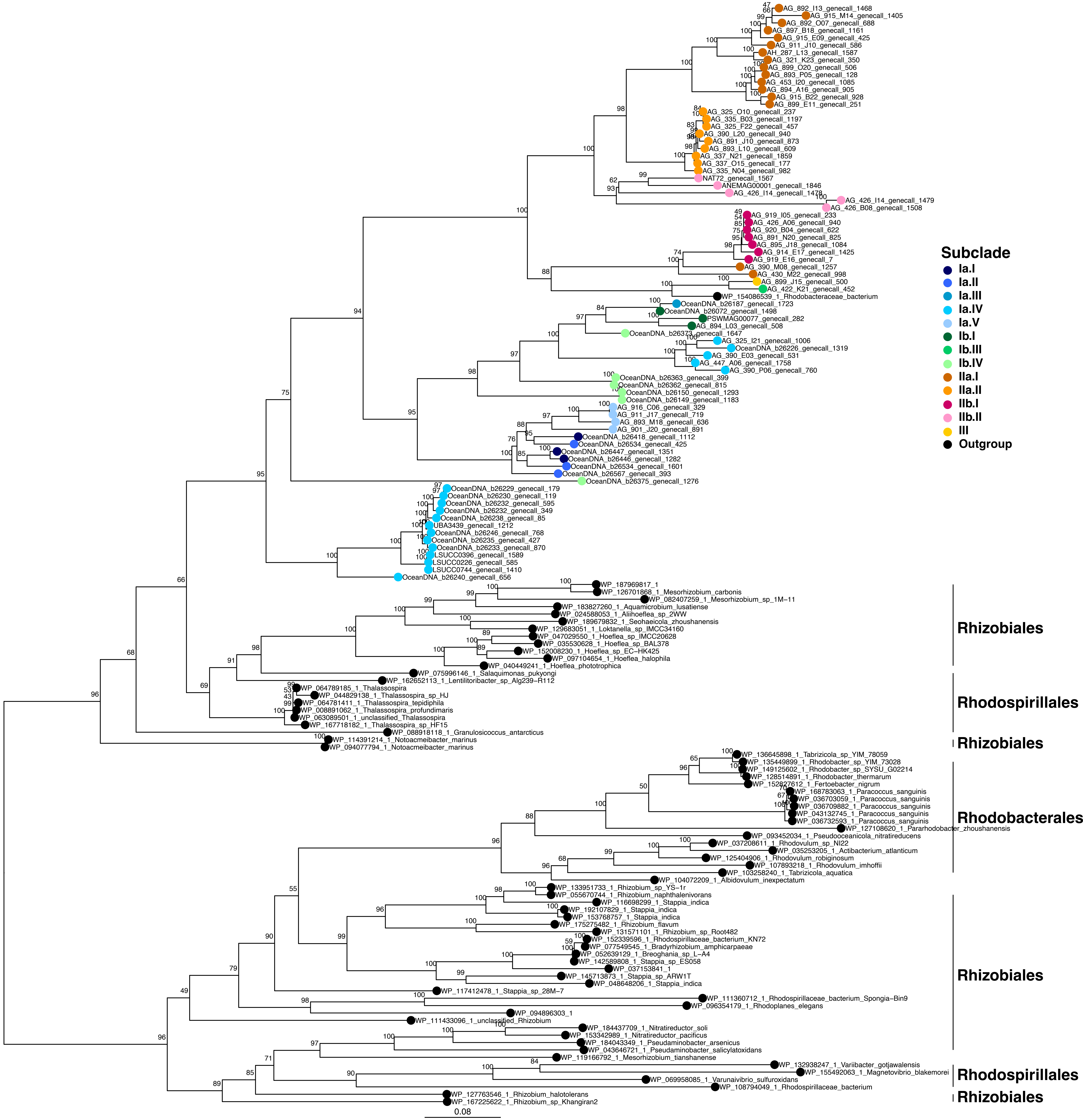

Subclade
Ia.I
Ia.II
Ia.III
Ia.IV
Ia.V
Ib.I
Ib.III
Ib.IV
IIa.I
IIa.II
IIb.I
IIb.II
III
Outgroup
AG_892_I13_genecall_1468
AG_915_M14_genecall_1405
AG_892_O07_genecall_688
AG_897_B18_genecall_1161
AG_915_E09_genecall_425
AG_911_J10_genecall_586
AH_287_L13_genecall_1587
AG_321_K23_genecall_350
AG_899_O20_genecall_506
AG_893_P05_genecall_128
AG_453_I20_genecall_1085
AG_894_A16_genecall_905
AG_915_B22_genecall_928
AG_899_E11_genecall_251
AG_325_O10_genecall_237
AG_335_B03_genecall_1197
AG_325_F22_genecall_457
AG_390_L20_genecall_940
AG_891_J10_genecall_873
AG_893_L10_genecall_609
AG_337_N21_genecall_1859
AG_337_O15_genecall_177
AG_335_N04_genecall_982
NAT72_genecall_1567
ANEMAG00001_genecall_1846
AG_426_I14_genecall_1478
AG_426_I14_genecall_1479
AG_426_B08_genecall_1508
AG_919_I05_genecall_233
AG_426_A06_genecall_940
AG_920_B04_genecall_622
AG_891_N20_genecall_825
AG_895_J18_genecall_1084
AG_914_E17_genecall_1425
AG_919_E16_genecall_7
AG_390_M08_genecall_1257
AG_430_M22_genecall_998
AG_899_J15_genecall_500
AG_422_K21_genecall_452
WP_154086539_1_Rhodobacteraceae_bacterium
OceanDNA_b26187_genecall_1723
OceanDNA_b26072_genecall_1498
PSWMAG00077_genecall_282
AG_894_L03_genecall_508
OceanDNA_b26373_genecall_1647
AG_325_I21_genecall_1006
OceanDNA_b26226_genecall_1319
AG_390_E03_genecall_531
AG_447_A06_genecall_1758
AG_390_P06_genecall_760
OceanDNA_b26363_genecall_399
OceanDNA_b26362_genecall_815
OceanDNA_b26150_genecall_1293
OceanDNA_b26149_genecall_1183
AG_916_C06_genecall_329
AG_911_J17_genecall_719
AG_893_M18_genecall_636
AG_901_J20_genecall_891
OceanDNA_b26418_genecall_1112
OceanDNA_b26534_genecall_425
OceanDNA_b26447_genecall_1351
OceanDNA_b26446_genecall_1282
OceanDNA_b26534_genecall_1601
OceanDNA_b26567_genecall_393
OceanDNA_b26375_genecall_1276
OceanDNA_b26229_genecall_179
OceanDNA_b26230_genecall_119
OceanDNA_b26232_genecall_595
OceanDNA_b26232_genecall_349
OceanDNA_b26238_genecall_85
UBA3439_genecall_1212
OceanDNA_b26246_genecall_768
OceanDNA_b26235_genecall_427
OceanDNA_b26233_genecall_870
LSUCC0396_genecall_1589
LSUCC0226_genecall_585
LSUCC0744_genecall_1410
OceanDNA_b26240_genecall_656
WP_187969817_1
WP_126701868_1_Mesorhizobium_carbonis
WP_082407259_1_Mesorhizobium_sp_1M−11
WP_183827260_1_Aquamicrobium_lusatiense
WP_024588053_1_Aliihoeflea_sp_2WW
WP_189679832_1_Seohaeicola_zhoushanensis
WP_129683051_1_Loktanella_sp_IMCC34160
WP_047029550_1_Hoeflea_sp_IMCC20628
WP_035530628_1_Hoeflea_sp_BAL378
WP_152008230_1_Hoeflea_sp_EC−HK425
WP_097104654_1_Hoeflea_halophila
WP_040449241_1_Hoeflea_phototrophica
WP_075996146_1_Salaquimonas_pukyongi
WP_162652113_1_Lentilitoribacter_sp_Alg239−R112
WP_064789185_1_Thalassospira
WP_044829138_1_Thalassospira_sp_HJ
WP_064781411_1_Thalassospira_tepidiphila
WP_008891062_1_Thalassospira_profundimaris
WP_063089501_1_unclassified_Thalassospira
WP_167718182_1_Thalassospira_sp_HF15
WP_088918118_1_Granulosicoccus_antarcticus
WP_114391214_1_Notoacmeibacter_marinus
WP_094077794_1_Notoacmeibacter_marinus
WP_136645898_1_Tabrizicola_sp_YIM_78059
WP_135449899_1_Rhodobacter_sp_YIM_73028
WP_149125602_1_Rhodobacter_sp_SYSU_G02214
WP_128514891_1_Rhodobacter_thermarum
WP_152827612_1_Fertoebacter_nigrum
WP_168783063_1_Paracoccus_sanguinis
WP_036703059_1_Paracoccus_sanguinis
WP_036709882_1_Paracoccus_sanguinis
WP_043132745_1_Paracoccus_sanguinis
WP_036732593_1_Paracoccus_sanguinis
WP_127108620_1_Pararhodobacter_zhoushanensis
WP_093452034_1_Pseudooceanicola_nitratireducens
WP_037208611_1_Rhodovulum_sp_NI22
WP_035253205_1_Actibacterium_atlanticum
WP_125404906_1_Rhodovulum_robiginosum
WP_107893218_1_Rhodovulum_imhoffii
WP_103258240_1_Tabrizicola_aquatica
WP_104072209_1_Albidovulum_inexpectatum
WP_133951733_1_Rhizobium_sp_YS−1r
WP_055670744_1_Rhizobium_naphthalenivorans
WP_116698299_1_Stappia_indica
WP_192107829_1_Stappia_indica
WP_153768757_1_Stappia_indica
WP_175275482_1_Rhizobium_flavum
WP_131571101_1_Rhizobium_sp_Root482
WP_152339596_1_Rhodospirillaceae_bacterium_KN72
WP_077549545_1_Bradyrhizobium_amphicarpaeae
WP_052639129_1_Breoghania_sp_L−A4
WP_142589808_1_Stappia_sp_ES058
WP_037153841_1
WP_145713873_1_Stappia_sp_ARW1T
WP_048648206_1_Stappia_indica
WP_117412478_1_Stappia_sp_28M−7
WP_111360712_1_Rhodospirillaceae_bacterium_Spongia−Bin9
WP_096354179_1_Rhodoplanes_elegans
WP_094896303_1
WP_111433096_1_unclassified_Rhizobium
WP_184437709_1_Nitratireductor_soli
WP_153342989_1_Nitratireductor_pacificus
WP_184043349_1_Pseudaminobacter_arsenicus
WP_043646721_1_Pseudaminobacter_salicylatoxidans
WP_119166792_1_Mesorhizobium_tianshanense
WP_132938247_1_Variibacter_gotjawalensis
WP_155492063_1_Magnetovibrio_blakemorei
WP_069958085_1_Varunaivibrio_sulfuroxidans
WP_108794049_1_Rhodospirillaceae_bacterium
WP_127763546_1_Rhizobium_halotolerans
WP_167225622_1_Rhizobium_sp_Khangiran2
Rhizobiales
Rhodospirillales
Rhizobiales
Rhodobacterales
Rhizobiales
Rhodospirillales
Rhizobiales
0.08

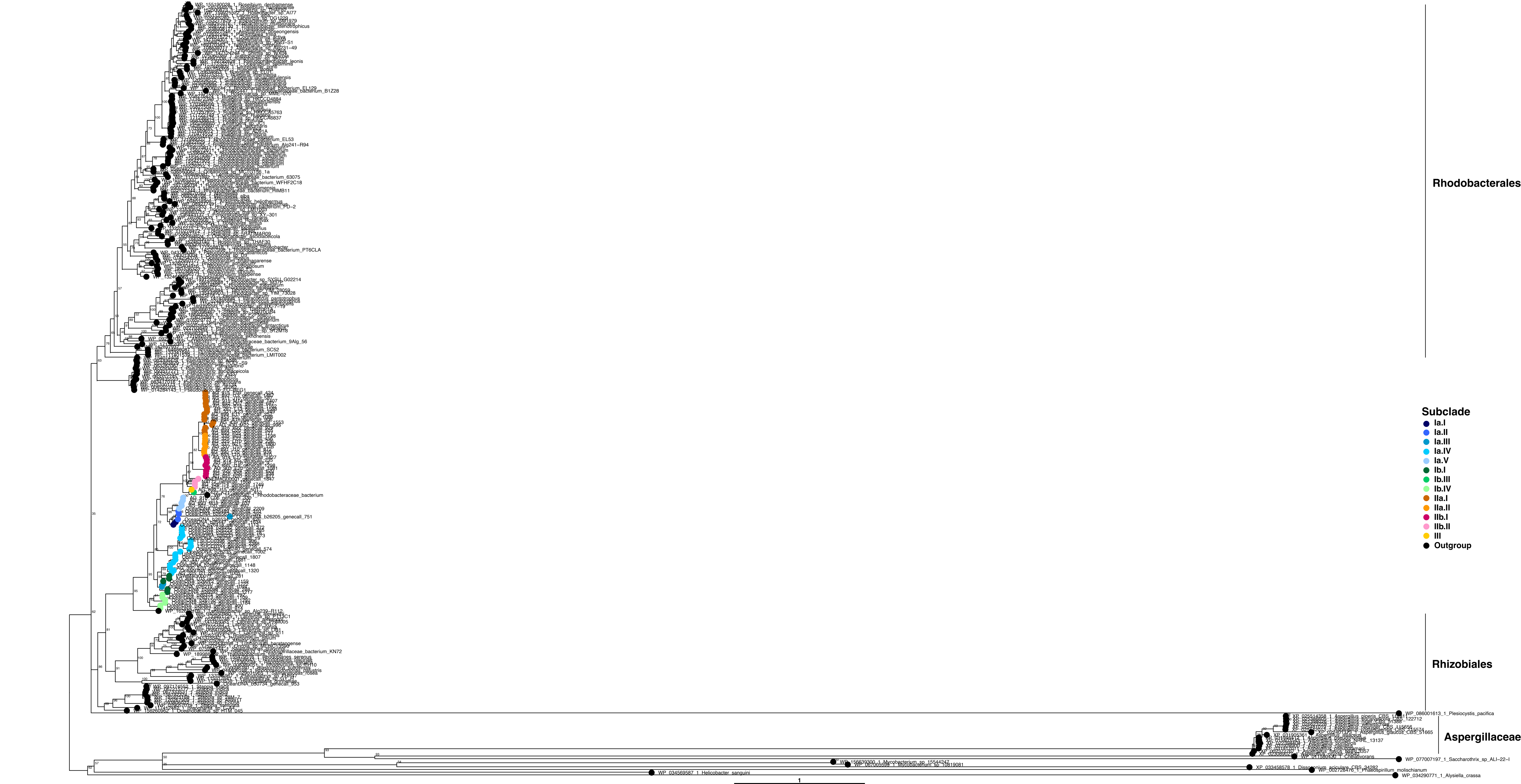

Rhodobacterales
Rhizobiales
Aspergillaceae
Subclade
Ia.I
Ia.II
Ia.III
Ia.IV
Ia.V
Ib.I
Ib.III
Ib.IV
IIa.I
IIa.II
IIb.I
IIb.II
III
Outgroup
WP_086001613_1_Plesiocystis_pacifica
WP_077007197_1_Saccharothrix_sp_ALI-22-I
WP_034569587_1_Helicobacter_sanguini
WP_034290771_1_Alysiella_crassa
1

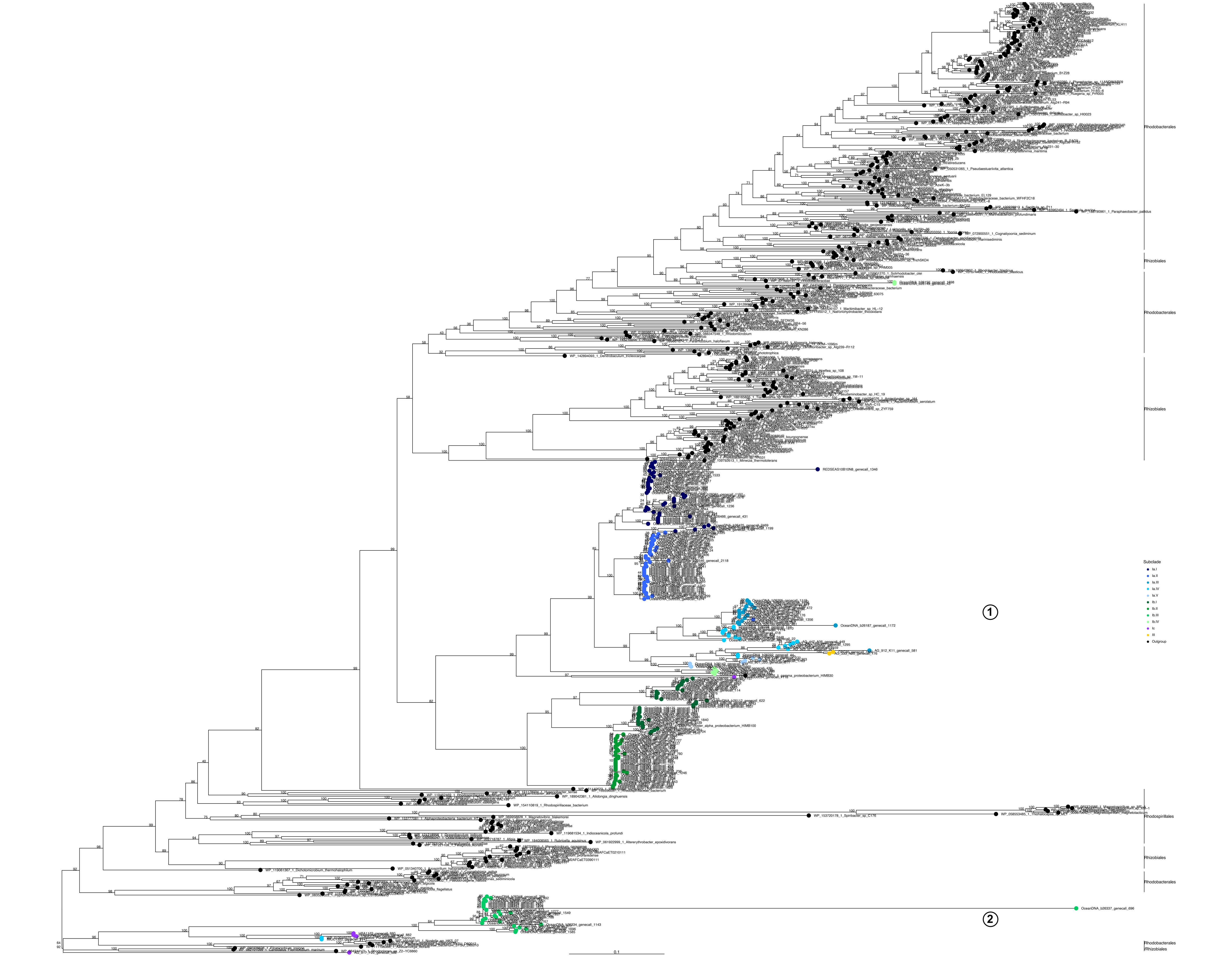
Rhodobacterales
Rhizobiales
Rhodobacterales
Rhizobiales
Subclade
Ia.I
Ia.II
Ia.III
Ia.IV
Ia.V
Ib.I
Ib.II
Ib.III
Ib.IV
Ic
III
Outgroup
1
Rhodospirillales
Rhizobiales
Rhodobacterales
2
Rhodobacterales
/Rhizobiales
0.1

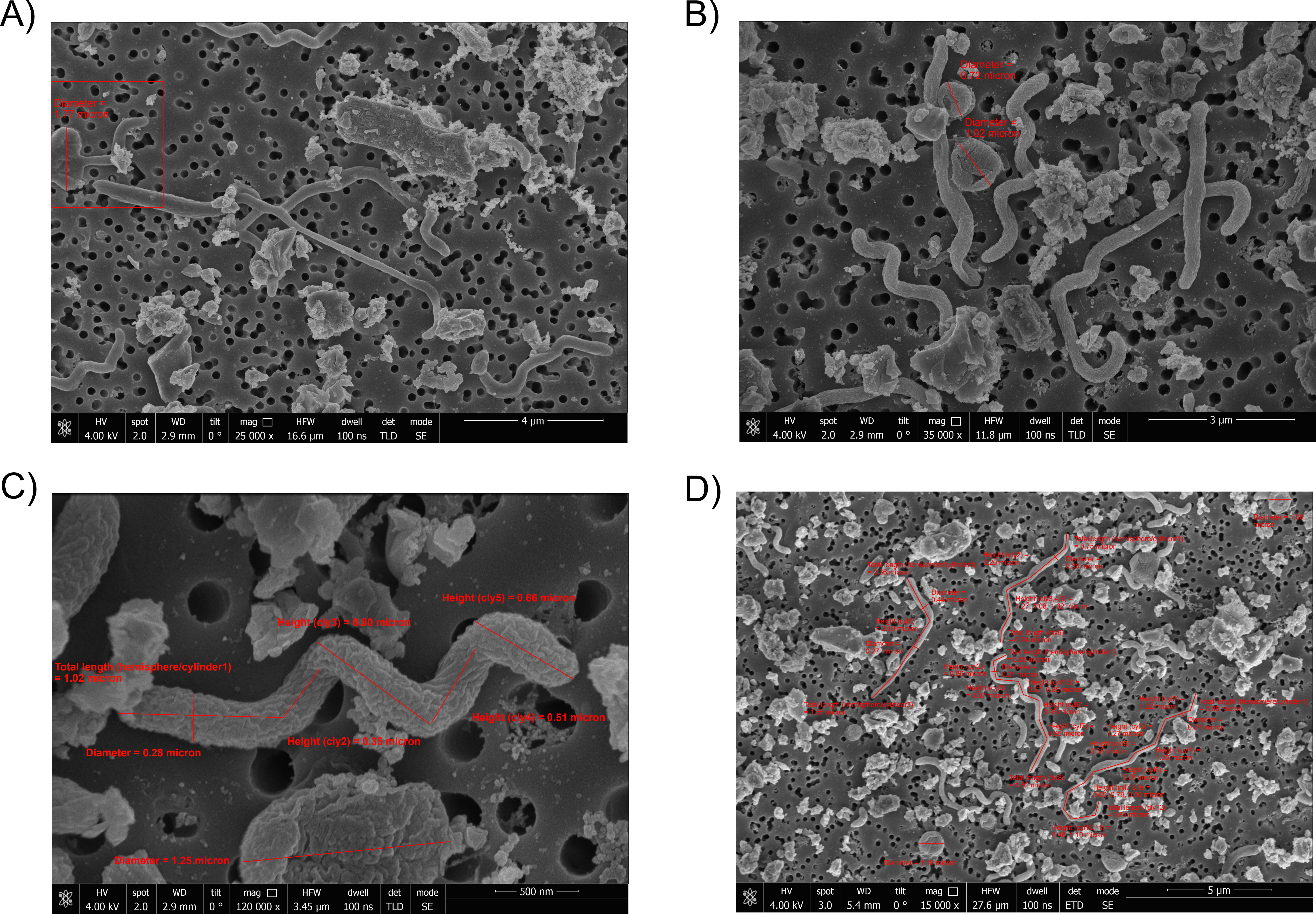
A)
Diameter = 1.77 micron
HV 4.00 kV spot 2.0 WD 2.9 mm tilt 0 ° mag 25 000 x HFW 16.6 µm dwell 100 ns det TLD mode SE
4 µm
B)
Diameter = 0.72 micron
Diameter = 1.02 micron
HV 4.00 kV spot 2.0 WD 2.9 mm tilt 0 ° mag 35 000 x HFW 11.8 µm dwell 100 ns det TLD mode SE
3 µm
C)
Height (cly5) = 0.66 micron
Height (cly3) = 0.80 micron
Total length (hemisphere/cylinder1) = 1.02 micron
Height (cly4) = 0.51 micron
Height (cly2) = 0.35 micron
Diameter = 0.28 micron
Diameter = 1.25 micron
HV 4.00 kV spot 2.0 WD 2.9 mm tilt 0 ° mag 120 000 x HFW 3.45 µm dwell 100 ns det TLD mode SE
500 nm
D)
HV 4.00 kV spot 3.0 WD 5.4 mm tilt 0 ° mag 15 000 x HFW 27.6 µm dwell 100 ns det ETD mode SE
5 µm
